# The type 2 diabetes gene product STARD10 is a phosphoinositide binding protein that controls insulin secretory granule biogenesis

**DOI:** 10.1101/2020.03.25.007286

**Authors:** Gaelle R. Carrat, Elizabeth Haythorne, Alejandra Tomas, Leena Haataja, Andreas Müller, Peter Arvan, Alexandra Piunti, Kaiying Cheng, Mutian Huang, Timothy J. Pullen, Eleni Georgiadou, Theodoros Stylianides, Nur Shabrina Amirruddin, Victoria Salem, Walter Distaso, Andrew Cakebread, Kate J. Heesom, Philip A. Lewis, David J. Hodson, Linford J. Briant, Annie C.H. Fung, Richard B. Sessions, Fabien Alpy, Alice P.S. Kong, Peter I. Benke, Federico Torta, Adrian Kee Keong Teo, Isabelle Leclerc, Michele Solimena, Dale B. Wigley, Guy A. Rutter

## Abstract

**Objective:** Risk alleles for type 2 diabetes at the *STARD10* locus are associated with lowered *STARD10* expression in the β-cell, impaired glucose-induced insulin secretion and decreased circulating proinsulin:insulin ratios. Although likely to serve as a mediator of intracellular lipid transfer, the identity of the transported lipids, and thus the pathways through which STARD10 regulates β-cell function, are not understood. The aim of this study was to identify the lipids transported and affected by STARD10 in the β-cell and its effect on proinsulin processing and insulin granule biogenesis and maturation.

**Methods:** We used isolated islets from mice deleted selectively in the β-cell for *Stard10* (β*StarD10*KO) and performed electron microscopy, pulse-chase, RNA sequencing and lipidomic analyses. Proteomic analysis of STARD10 binding partners was executed in INS1 (832/13) cell line. X-ray crystallography followed by molecular docking and lipid overlay assay were performed on purified STARD10 protein.

**Results:** β*StarD10*KO islets had a sharply altered dense core granule appearance, with a dramatic increase in the number of “rod-like” dense cores. Correspondingly, basal secretion of proinsulin was increased. Amongst the differentially expressed genes in β*StarD10*KO islets, expression of the phosphoinositide binding proteins *Pirt* and *Synaptotagmin 1* were decreased while lipidomic analysis demonstrated changes in phosphatidyl inositol levels. The inositol lipid kinase PIP4K2C was also identified as a STARD10 binding partner. STARD10 bound to inositides phosphorylated at the 3’ position and solution of the crystal structure of STARD10 to 2.3 Å resolution revealed a binding pocket capable of accommodating polyphosphoinositides.

**Conclusion:** Our data indicate that STARD10 binds to, and may transport, phosphatidylinositides, influencing membrane lipid composition, insulin granule biosynthesis and insulin processing.

## 1. Introduction

Diabetes mellitus is characterised by high blood glucose and currently affects around 8.5% of the population worldwide [1]. Normal glucose homeostasis requires the processing of proinsulin and the storage of the mature hormone within dense core granules in the pancreatic β-cell [2]. Glucose-induced insulin secretion involves glucose uptake and metabolism through the glycolytic pathway, increased ATP production by the mitochondria and the closure of ATP-sensitive K^+^ channels (K_ATP_). Subsequent depolarization of the plasma membrane leads to opening of voltage-gated Ca^2+^ channels, Ca^2+^ dependent assembly of the SNARE (SNAP (Soluble NSF Attachment Protein) Receptor) complex and exocytosis [3]. In the intact islet, β-cell-β-cell connections allow coordinated insulin secretion through propagation of Ca^2+^ and other signals [4,5,6] a process impaired by glucolipotoxicity [7], and affected by genes implicated in diabetes risk through genome-wide association studies (GWAS) such as *ADCY5* [8] and *TCF7L2* [9].

We have recently examined a T2D-associated locus adjacent to *STARD10* on chromosome 11q13 [10,11]. Risk variants at this locus were associated with a decrease in *STARD10* mRNA in human islets, with no concomitant change in the liver. Changes in the expression of the nearby *ARAP1* gene were not associated with the possession of risk alleles in either tissue, pointing to STARD10 as the mediator of the effects of risk variants. Providing further compelling evidence for *STARD10* as an “effector” gene, mice deleted for *Stard10* specifically in the β-cell recapitulated the features observed in the human carriers of the risk allele, with an increase in fed glycemia and a decrease in the plasma proinsulin:insulin ratio. Islets isolated from the knock-out mice also showed impaired glucose-induced Ca^2+^ signalling and insulin secretion. Thus, β-cell STARD10 may be a useful therapeutic target in some forms of type 2 diabetes, particularly in risk allele carriers who may benefit from a tailored, pharmacogenetic approach.

STARD10 (previously named phosphatidylcholine transfer protein-like, Pctp-l) is a phospholipid transfer protein possessing a steroidogenic acute regulatory protein (StAR)-related lipid transfer (StART) domain that facilitates the transport of phosphatidylcholine and phosphatidylethanolamine between intracellular membranes [12]. Nevertheless, the molecular mechanisms by which STARD10 regulates insulin secretion in the β-cell, as well as its subcellular localisation and target membranes, remain unknown. We therefore examined in detail here the role of STARD10 in controlling the lipid composition, granule maturation, proinsulin processing and metal ion homeostasis in the mouse β-cell. We reveal an unexpected role for STARD10 in binding inositol phospholipids which may contribute to both secretory granule biogenesis and intracellular signalling.

## 2. Material and methods

### 2.1. Generation and use of Stard10 null mice

All animal procedures were approved by the U.K. Home Office according to the Animals (Scientific Procedures) Act 1986 of the United Kingdom (PPL PA03F7F0F to I.L.). *Stard10* whole body and conditional KO mice (C57BL/6NTac background) were generated by the trans-NIH Knock-Out Mouse Project (KOMP) and obtained from the KOMP Repository via the international mouse phenotyping consortium (IMPC). Mice homozygous for floxed *Stard10* (*Stard10*^fl/fl^) alleles were crossed to mice expressing *Cre* recombinase from the endogenous *Ins1* locus (*Ins1-Cre* mice). This generated *Stard10*^fl/fl^:*Ins1Cre*^+^ (β*Stard10*KO) mice as in [10].

### 2.2. Islet isolation and culture

Mice were euthanized by cervical dislocation and pancreatic islets isolated by collagenase digestion as previously described [13] and cultured in RPMI 1640 medium, 11 mM glucose, supplemented with 10% (v/v) fetal bovine serum plus penicillin (100 units/mL) and streptomycin (0.1 mg/mL) at 37°C in an atmosphere of humidified air (95%) and CO_2_ (5%).

### 2.3. Transmission electron microscopy imaging

For conventional EM, islets were chemically fixed in 2% paraformaldehyde (EM grade), 2% glutaraldehyde and 3 mM CaCl_2_ in 0.1 M cacodylate buffer for 2 h at room temperature then left overnight at 4°C in fresh fixative solution, osmicated, enrobed in agarose plugs, dehydrated in ethanol and embedded on Epon. Epon was polymerised overnight at 60°C. Ultrathin 70 nm sections were cut with a diamond knife (DiATOME) in a Leica Ultracut UCT ultramicrotome before examination on an FEI Tecnai G2 Spirit TEM. Images were acquired in a charge-coupled device camera (Eagle), and processed in ImageJ.

### 2.4. Measurements of islet Zn^2+^ concentrations

#### 2.4.1. Cytosolic free Zn^2+^ measurements

Imaging of cytosolic [Zn^2+^] using the eCALWY4 sensor was carried out on mouse islets dispersed onto coverslips as previously described [14]. Cells were maintained at 37°C and Krebs-HEPES-bicarbonate buffer (11 mM) was perifused with additions as stated in the figure. Images were captured at 433 nm monochromatic excitation wavelength. Acquisition rate was 20 images/min.

Image analysis was performed with ImageJ software [15] using a homemade macro and the fluorescence emission ratios were derived after subtracting background. We observed that during acquisition, photobleaching gradually decreased the steady-state ratio with a linear kinetic (not shown). This drift was thus, when necessary, corrected in function of time with a constant factor.

#### 2.4.2. Measurement of islet zinc content by inductively coupled plasma mass spectrometry (ICP-MS)

Mouse islets were washed twice in PBS and stored at −80°C until ready to process. Islets were lysed in 100 μL nitric acid (trace metal grade) and heated at 50°C for 6 hours. The samples were then cooled, diluted in trace metal grade water up to a final volume of 1.3 mL and 10 μL of 1000 ppb mix of 5 internal standards (bismuth, indium, scandium, terbium, and yttrium) was added. Standards between 0.5 and 500.5 μg/L of Zn were used for calibration. Samples were run on the Perkin Elmer NexION 350D using the Syngistix software.

### 2.5. Metabolic labelling of mouse pancreatic islets

Islets isolated from β*Stard10*KO and WT littermate mice were recovered overnight in RPMI-1640 medium containing 11 mM glucose plus 10% FBS and penicillin-streptomycin. In each case, 50 islets were washed twice in prewarmed RPMI lacking cysteine and methionine. Islets were pulse-labeled with ^35^S-labeled amino acids for 20 min and chased for 1.5 h or 4 h in 5 mM glucose RPMI-1640 plus 10% FBS, Hepes, Pyruvate and penicillin-streptomycin. Islets were lysed in radioimmunoprecipitation assay buffer (25 mmol/L Tris, pH 7.5; 1% Triton X-100; 0.2% deoxycholic acid; 0.1% SDS; 10 mmol/L EDTA; and 100 mmol/L NaCl) plus 2 mmol/L N-ethylmaleimide and protease inhibitor cocktail. Both cell lysates and media were precleared with pansorbin and immunoprecipitated with anti-insulin antibodies and protein A agarose overnight at 4°C. Immunoprecipitates were analyzed using tris-tricine-urea-SDS-PAGE under nonreducing conditions or SDS-PAGE on 4-12% acrylamide gradient gels (NuPAGE) under reducing conditions as indicated with phosphorimaging. Bands were quantified using ImageJ software.

### 2.6. In vitro proinsulin and insulin measurements

Islets (10/well) were incubated in triplicate for each condition and treatment. Islets were preincubated for 1 h in 3 mM glucose Krebs-Ringer-Hepes-Bicarbonate (KRH) buffer prior to secretion assay (30 min.) in 3 mM or 17 mM glucose. The secretion medium were then collected to measure the insulin and proinsulin concentrations using respectively an insulin HTRF kit (Cisbio Bioassays) and a rat/mouse proinsulin ELISA kit (Mercodia).

### 2.7. Lipidomic analysis

Islets isolated from β*Stard10*KO and WT littermate mice were recovered overnight in RPMI-1640 medium containing 11 mM glucose plus 10% FBS and penicillin-streptomycin. Islets were then washed twice in PBS, snap-frozen in a bath of ethanol and dry ice and kept at −80°C until ready to process. Lipids were extracted with 100 μl 1-butanol/methanol (1:1, v/v) containing 2.5 μL of SPLASH^™^ Lipidomix^®^ Mass Spec Standard I and 2.5 μL Cer/Sph Mixture I, purchased from Avanti Polar Lipids. The mixture was vortexed for 30 s, sonicated for 30 min. at 20°C and then centrifuged at 14,000 g for 10 min. The supernatant was transferred into vials. The lipidomic analysis was performed by the Singapore Lipidomics Incubator (SLING) using an Agilent 1290 Infinity II LC system combined with an Agilent 6495 triple quadrupole mass spectrometer. Reverse Phase chromatographic separation of 1 μL samples were carried out on an Agilent Zorbax RRHD Eclipse Plus C18, 95 Å (50 x 2.1 mm, 1.8 μm) column maintained at 40°C. The mobile phases consisted of (A) 10 mmol/L ammonium formate in acetonitrile/water (40:60, v/v) and (B) 10 mmol/L ammonium formate in acetonitrile/2-propanol (10:90, v/v). Using a flow rate of 0.4 mL/min., the gradient elution program included 20% B to 60% B from 0 to 5 min., 60 to 100% B from 2 to 7 min, where it was maintained till 9 min., then reequilibrated at 20% B for 1.8 min. prior to the next injection. All samples were kept at 10°C in the autosampler. The lipid amounts were normalised to protein content. MRM chromatograms obtained in positive ion mode, and covering >10 lipid classes, were processed using Agilent MassHunter Quantitative Analysis software (version B.08.00). Peaks were annotated based on retention time and specific MRM transitions.

### 2.8. Massive parallel sequencing of RNA (RNAseq)

Total RNA was extracted with TRIzol from isolated mouse islets. Polyadenylated transcripts were selected during the preparation of paired-end, directional RNAseq libraries using the Illumina TruSeq Stranded mRNA Library Prep Kit. Libraries were sequenced on an Illumina HiSeq 4000 machine at 75 bp paired end. The quality of the sequenced libraries was assessed using fastQC. Reads were mapped to the Grc38m assembly using HiSat2. Annotated transcripts were quantified using featureCounts, and differentially expressed genes were identified with DESeq2. Raw sequence data for RNAseq will be made available via deposition to ArrayExpress.

### 2.9. Purification and identification of STARD10 interacting proteins by mass spectrometry

#### 2.9.1. Co-immunoprecipitation

INS1 (832/13) cells were lysed in the following non-denaturing lysis buffer: 20 mM HEPES, 150 mM NaCl, 1% Igepal, protease inhibitors (Roche Diagnostics, complete, EDTA-free protease inhibitor cocktail tablets), phosphatase inhibitors (Sigma, P5726). 6 μg of Rabbit IgG isotype control (Abcam, ab171870) or anti-PCTP-L (STARD10) antibody (Abcam. ab242109) were bound to 50 μL of Dynabeads protein A for 1 h at 4°C. For co-immunoprecipitation (Co-IP), 1 mg of protein lysate was incubated with the complex beads-antibodies overnight at 4°C. Beads were then washed twice in lysis buffer and twice in PBS-Tween 0.01% prior to proteomic analysis by the Bristol proteomics facility.

#### 2.9.2. TMT Labelling and High pH reversed-phase chromatography

Pull-down samples were reduced (10 mM TCEP 55°C, 1 h), alkylated (18.75 mM iodoacetamide, room temperature, 30 min.) and digested on the beads with trypsin (2.5 μg trypsin; 37°C, overnight), then labelled with Tandem Mass Tag (TMT) six plex reagents according to the manufacturer’s protocol (Thermo Fisher Scientific, Loughborough, LE11 5RG, UK) and the labelled samples pooled.

The pooled sample was evaporated to dryness, resuspended in 5% formic acid and then desalted using a SepPak cartridge according to the manufacturer’s instructions (Waters, Milford, Massachusetts, USA). Eluate from the SepPak cartridge was again evaporated to dryness and resuspended in buffer A (20 mM ammonium hydroxide, pH 10) prior to fractionation by high pH reversed-phase chromatography using an Ultimate 3000 liquid chromatography system (Thermo Fisher Scientific). In brief, the sample was loaded onto an XBridge BEH C18 Column (130 Å, 3.5 μm, 2.1 mm x 150 mm, Waters, UK) in buffer A and peptides eluted with an increasing gradient of buffer B (20 mM Ammonium Hydroxide in acetonitrile, pH 10) from 0-95% over 60 minutes. The resulting fractions (4 in total) were evaporated to dryness and resuspended in 1% formic acid prior to analysis by nano-LC MSMS using an Orbitrap Fusion Tribrid mass spectrometer (Thermo Scientific).

#### 2.9.3. Nano-LC Mass Spectrometry

High pH RP fractions were further fractionated using an Ultimate 3000 nano-LC system in line with an Orbitrap Fusion Tribrid mass spectrometer (Thermo Scientific). In brief, peptides in 1% (vol/vol) formic acid were injected onto an Acclaim PepMap C18 nano-trap column (Thermo Scientific). After washing with 0.5% (vol/vol) acetonitrile 0.1% (vol/vol) formic acid peptides were resolved on a 250 mm × 75 μm Acclaim PepMap C18 reverse phase analytical column (Thermo Scientific) over a 150 min. organic gradient, using 7 gradient segments (1-6% solvent B over 1 min., 6-15% B over 58 min., 15-32%B over 58 min., 32-40%B over 5 min., 40-90%B over 1 min., held at 90%B for 6 min. and then reduced to 1%B over 1 min.) with a flow rate of 300 nl min^−1^. Solvent A was 0.1% formic acid and Solvent B was aqueous 80% acetonitrile in 0.1% formic acid. Peptides were ionized by nano-electrospray ionization at 2.0 kV using a stainless steel emitter with an internal diameter of 30 μm (Thermo Scientific) and a capillary temperature of 275°C.

All spectra were acquired using an Orbitrap Fusion Tribrid mass spectrometer controlled by Xcalibur 3.0 software (Thermo Scientific) and operated in data-dependent acquisition mode using an SPS-MS3 workflow. FTMS1 spectra were collected at a resolution of 120 000, with an automatic gain control (AGC) target of 400 000 and a max injection time of 100 ms. Precursors were filtered with an intensity range from 5000 to 1E20, according to charge state (to include charge states 2-6) and with monoisotopic peak determination set to peptide. Previously interrogated precursors were excluded using a dynamic window (60 s ± 10 ppm). The MS2 precursors were isolated with a quadrupole isolation window of 1.2m/z. ITMS2 spectra were collected with an AGC target of 10 000, max injection time of 70ms and CID collision energy of 35%.

For FTMS3 analysis, the Orbitrap was operated at 30 000 resolution with an AGC target of 50 000 and a max injection time of 105 ms. Precursors were fragmented by high energy collision dissociation (HCD) at a normalised collision energy of 55% to ensure maximal TMT reporter ion yield. Synchronous Precursor Selection (SPS) was enabled to include up to 5 MS2 fragment ions in the FTMS3 scan.

#### 2.9.4. Data Analysis

The raw data files were processed and quantified using Proteome Discoverer software v2.1 (Thermo Scientific) and searched against the UniProt Rat database (downloaded January 2019; 35759 entries) using the SEQUEST algorithm. Peptide precursor mass tolerance was set at 10 ppm, and MS/MS tolerance was set at 0.6 Da. Search criteria included oxidation of methionine (+15.9949) as a variable modification and carbamidomethylation of cysteine (+57.0214) and the addition of the TMT mass tag (+229.163) to peptide N-termini and lysine as fixed modifications. Searches were performed with full tryptic digestion and a maximum of 2 missed cleavages were allowed. The reverse database search option was enabled and all data was filtered to satisfy false discovery rate (FDR) of 5%.

### 2.10. Lipid overlay assay

All incubation steps were performed at room temperature. PIP strips (Thermo Scientific) were blocked for 1h in TBS containing 0.1% Tween-20 (TBS-T) supplemented by 3% fatty-acid free BSA (Sigma Aldrich) before incubation with the purified STARD10 protein (1 μg/mL in TBS-T + 3% BSA) for 1 h. The membrane was washed 5 times 10 min. in TBS-T and probed for 1 h with the polyclonal goat anti-STARD10 antibody (Santa Cruz sc54336; 1/1000 in TBS-T + 3%BSA). After 5 washes of 10 min. in TBS-T, the membrane was incubated 1 h with the horseradish peroxidase conjugated donkey anti-goat IgG (Santa Cruz sc2020; 1/2000 in TBS-T + 3%BSA). After 5 washes of 10 min. in TBS-T, bound proteins were detected by ECL reagent (GE Healthcare).

### 2.11. Structure solution of STARD10

#### 2.11.1. Cloning

The full length gene encoding human STARD10 protein was amplified from cDNA by PCR and cloned into a modified pET28a expression vector, pET28-HMT, which contains a fused N-terminal 6×His-tag, an MBP-tag and a TEV protease recognition site (His-MBP-TEV) by Infusion^®^ HD Cloning kit (Takara Bio, USA). The fidelity of the constructs was confirmed by gel electrophoresis and sequencing.

#### 2.11.2. Protein preparation

In brief, transformed E. Coli BL21 (DE3) clones were grown at 37°C in LB medium containing 50 μg/mL Kanamycin to an optical density at 600 nm of 0.8. Protein expression was induced at 30°C for 4 h by adding isopropyl-β-D-thiogalactopyranoside (IPTG) to a final concentration of 0.5 mM. After harvesting, cells were re-suspended in lysis buffer (20 mM Tris (pH 8.0), 1 M NaCl and 0.5 mM TCEP) with protease inhibitor, lysed by sonication and centrifuged at 18,000 × g for 60 min. at 4°C. The supernatant was loaded on a HisTrap HP column (GE Healthcare, Fairfield, CT), equilibrated with buffer A (20 mM Tris (pH 8.0), 1 M NaCl, 0.5 mM TCEP and 5 mM imidazole), washed with 30 mM imidazole and finally eluted with 500 mM imidazole. After His-MBP-TEV-tag-removal using TEV protease, the protein was dialysed into buffer B (20 mM Tris (pH 8.0), 100 mM NaCl and 0.5 mM TCEP) and re-loaded onto the HisTrap HP column (GE Healthcare) to remove the tag, uncleaved protein and TEV protease. The flowthrough fractions were collected and loaded onto a MonoQ column (GE Healthcare) preequilibrated with buffer B. Some *E.coli* background protein and DNA rather than STARD10 can bind on the MonoQ. Then the flow-through fractions were collected and loaded onto a Heparin HP column (GE Healthcare) pre-equilibrated with buffer B. Fraction containing STARD10 protein was eluted with a linear gradient from 250 mM to 800 mM NaCl. The protein was finally purified by Superdex 75 10/300 GL column (GE Healthcare) with buffer B.

#### 2.11.3. Crystallization and structure determination

Crystallization trials were carried out by the sitting drop vapor diffusion method at 293 K. Freshly purified STARD10 was concentrated to ~38 mg/mL and centrifuged to remove insoluble material before crystallization. Single crystals appeared in condition containing 50% (v/v) PEG 200, 100 mM Sodium phosphate dibasic/Potassium phosphate monobasic (pH 6.2) and 200 mM NaCl after one month. Cryo-freezing was achieved by stepwise soaking the crystals in reservoir solution containing 10, 20, and 30% (w/v) glycerol for 3 min. and flash freezing in liquid nitrogen. X-ray diffraction data were collected on beamline I03 at the Diamond synchrotron X-ray source, and were integrated and scaled with the Xia2 system [16].

Structure was determined by molecular replacement using the START domain structure from human STARD5 protein (PDB ID: 2R55) as search model in CCP4, followed by rigid-body refinement by Refmac5 [17]. Structure was refined using PHENIX [18] and interspersed with manual model building using COOT [19]. The structure contains one STARD10 molecule in the asymmetric unit. 245 out of 291 residues (STARD10 full length) were successfully built into the density in the final structure. The statistics for data collection and model refinement are listed in Supplemental Table 6.

### 2.12. Molecular docking of ligands in STARD10 and STARD2

In silico molecular docking was performed using the Bristol University Docking Engine (BUDE 1.2.9). Firstly, a set of conformations for the head groups, glycero-inositol-1-phosphate and glycero-3-phosphoinositol-1-phosphate were generated using OpenBabel (2.4.1) giving 87 and 85 conformers respectively. The docking grid was centered on the central cavity in STARD2 (1LNL) and STARD10 (6SER). Each conformer of the PI and PI(3)P headgroups was docked into both structures. Each docking run found the best solutions by sampling a total of 1.1 million poses via BUDE’s genetic algorithm. Final models were constructed by adding the 2 linoleoyl chains from the conformation in 1LN1 (STARD2 dilinoleoylphosphatidylcholine complex) and refined by energy minimisation with GROMACS.

### 2.13. Statistics

Data are expressed as mean ± SD. Normality of the data distribution was tested by D’Agostino and Pearson and Shapiro-Wilk normality tests. For normally distributed data, significance was tested by Student’s two-tailed t-test or Welch’s t-test if the variances were found significantly different by *F*-test. Mann-Whitney test was used for non-parametric data, and one-or two-way ANOVA with SIDAK multiple comparison test for comparison of more than two groups, using Graphpad Prism 7 software. *p* < 0.05 was considered significant.

## 3. Results

### 3.1. Stard10 deletion affects dense core granule ultrastructure

As an initial approach to determining the target membranes for STARD10 action, we first explored the impact of deleting the *Stard10* gene on β-cell ultrastructure. We have previously shown that crossing of *Stard10* floxed mice to Ins1Cre knock-in mice [20] efficiently and selectively deletes STARD10 in the pancreatic β-cell [10]. Transmission electron microscopy images of β-cells in islets isolated from β*Stard10*KO mice revealed a dramatic change in insulin granule morphology with a significant increase in “atypical” granules with a “rod-shaped” core (Figure 1A and B; *Cre*+: 12.05 ± 1.67% *versus Cre*-: 2.78 ± 0.36%; *p*<0.001, Student’s *t* test, *n* = 3 animals). In addition, the mean granule diameter was decreased (Figure 1C; *Cre+:* 255.2 ± 25.8 nm *versus Cre*-: 277.9 ± 24.9 nm; *p*<0.001, Student’s *t*-test, *n* = 42 images from 3 animals) and the “circularity” increased slightly (Figure 1D; *Cre*+: 0.89 ± 0.01 *versus Cre*-: 0.87 ± 0.03; *p*<0.05, Welch’s *t*-test, *n* = 42 images from 3 animals) in β*Stard10*KO compared to WT littermate β-cells. However, the cytoplasmic abundance of granule profiles (“density”) (Figure 1E; *Cre*-: 3.82 ± 1.80 *vs. Cre*+: 3.66 ± 0.61 granules/μm^2^; ns, Mann-Whitney test, *n* = 19 images from 3 animals) and the number of granules morphologically docked to the plasma membrane (Figure 1F; *Cre*-: 0.76 ± 0.43 *vs. Cre*+: 0.75 ± 0.28 granules/μm plasma membrane; ns, Mann-Whitney test, *n* = 18 images from 3 animals; granules were considered morphologically docked if the distance from their centre to the plasma membrane was ≤ 200nm) remained similar in both genotypes.

**Figure 1:**
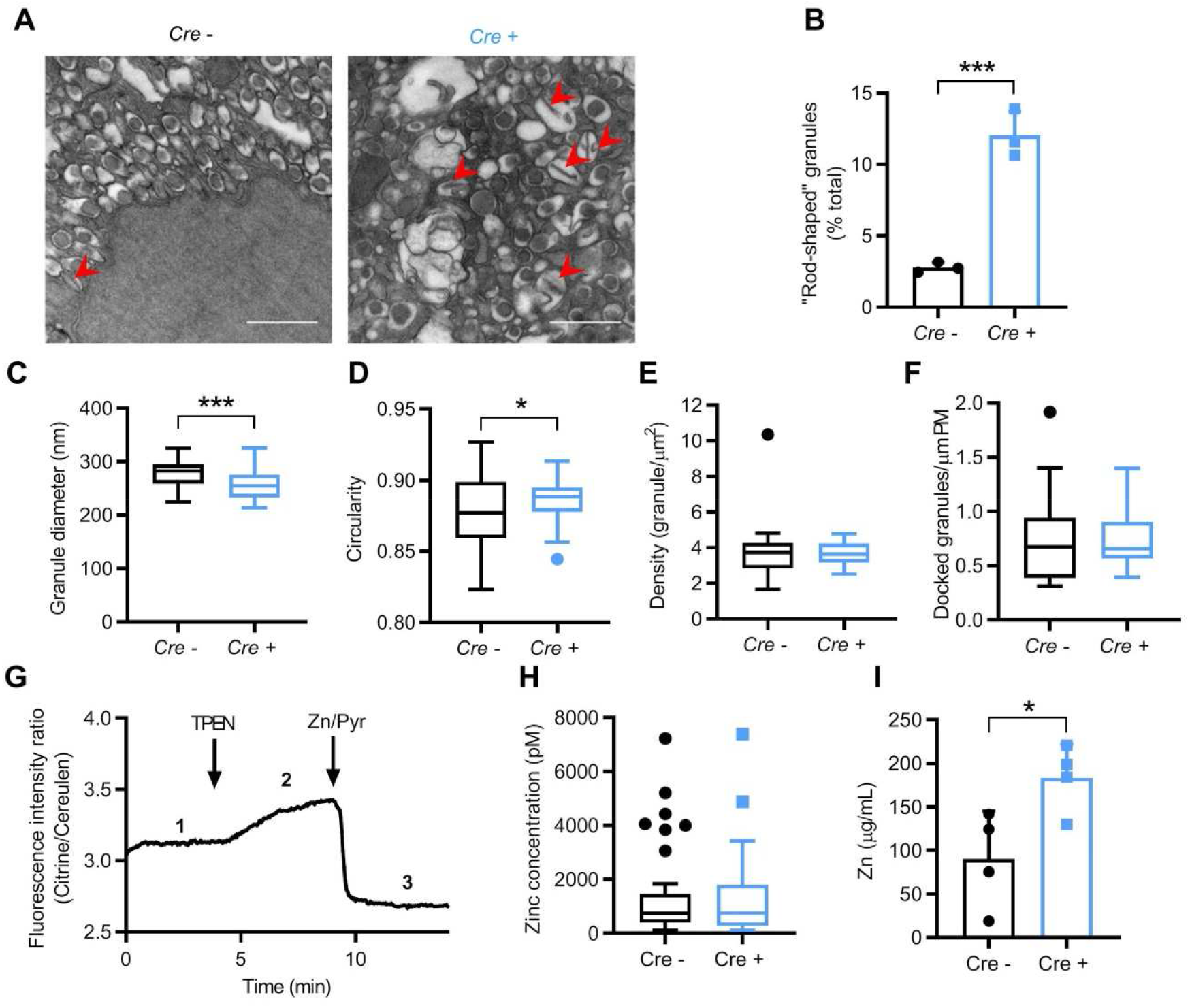
βStard10KO β-cells display altered granule morphology. A, Representative Transmission Electron Microscopy images of *Cre*- and *Cre*+ β-cells. Red arrowhead: granules with a “rod-shaped” core. Scale bar = 1μm. B, “Rod-shaped” core granule numbers are increased in the β*Stard10*KO β-cells (*n* = 3 animals, 6 images per animal; *p*<0.001, Student’s *t*-test). C, β-cell granule diameter (nm) (*n* = 42 images from 3 animals, *p*<0.001, Student’s *t*-test). D, β-cell granule “circularity” (*n* = 42 images from 3 animals, *p*<0.05, Welch’s *t*-test). E, β-cell granule density (*n* = 19 images from 3 animals, ns, Mann-Whitney test). F, β-cell morphologically docked granules (per μm plasma membrane) (*n* = 18 images from 3 animals, ns, Mann-Whitney test). G, Representative trace for β-cell expressing the cytosolic eCALWY4 Zn^2+^ sensor. Steady state fluorescence intensity ratio (citrine/cerulean) (1, R) was first measured before the maximum ratio (2, Rmax) was obtained under perfusion with buffer containing 50 μM TPEN (zinc-free condition). Finally, the minimum ratio (3, Rmin) was obtained under perfusion with buffer containing 5 μM pyrithione and 100 μM Zn^2+^ (zinc-saturated condition). Cytosolic free Zn^2+^ concentrations were calculated using the formula: (R-Rmin)/(Rmax-Rmin). H, Cytosolic Zn^2+^ concentrations measured by eCALWY4 in *Cre*- and *Cre*+ β-cells (n = 33 – 65 cells per genotype, ns, Mann-Whitney test). I, Total islet zinc measured by inductively coupled plasma mass spectrometry (ICP/MS) in *Cre*- and *Cre*+ animals (n = 4 animals/genotype, *p<0.05, unpaired t-test)

Analyses of transmission electron microscopy images of human β-cells from partially pancreatectomised patient samples (Supplemental Figure 1A and Supplemental Table 1) showed no significant correlation between *STARD10* expression measured by RNAseq and the percentage of mature granules (Supplemental Figure 1B) or the density of mature, immature or total granules (Supplemental Figure 1C).

In conclusion, deleting *Stard10* specifically in the β-cell in mice greatly affected insulin granule ultrastructure and overall shape with an increase in granules with a “rod-shaped” core.

### 3.2. Islet zinc content is increased in βStard10KO mice

To determine whether the abnormalities in granule structure may result from altered Zn^2+^ content of granules, itself a critical regulator of insulin crystallisation [2,21], we used the Förster resonance energy transfer- (FRET) based sensor eCALWY4 [14] to measure free Zn^2+^ concentration in the cytosol of dissociated islets from β*Stard10*KO mice. The measured free cytosolic Zn^2+^ concentrations were similar in both genotypes (Figure 1G and H; ns; Mann Whitney test, n = 33–65 cells). On the other hand, total zinc, measured by inductively-coupled plasma mass spectrometry (ICP/MS) was higher in β*Stard10*KO isolated islets compared to WT ones (Figure 1I: *Cre*-: 90.1 ± 55.15 *vs. Cre*+: 183.4 ± 38.73 μg/mL; *p*<0.05; unpaired t test, n = 4 animals).

### 3.3. Newly synthesised proinsulin is constitutively secreted by βStard10KO islets

The characteristic decrease of plasma proinsulin:insulin ratio observed in human carriers of the risk alleles at this locus [22], and the observation of a similar phenotype in β*Stard10*KO mice [10], suggest an action of STARD10 on proinsulin processing in the β-cell. We therefore next investigated this hypothesis by performing a metabolic labelling pulse-chase experiment in isolated islets from WT and β*Stard10*KO mice.

After 20 min. of pulse-labelling with ^35^S-amino acids, the islets of both genotypes were chased for 1.5 or 4 h in 5.5 mM glucose medium. Insulin was immunoprecipitated from both the cell lysate (C) and the secretion media (M) (Figure 2A). Labeled proinsulin secretion by *Cre*+ islets was increased at low (5.5 mM) glucose compared to *Cre*-mice after 4 h of chase (Figure 2B; *Cre*-: 8.46 ± 2.11 *vs. Cre*+: 12.64 ± 2.42% total; \**p*<0.05, Mann Whitney test, *n* = 4-5 animals). This increase in basal proinsulin secretion in the KO islets tended to be observed from 1.5 h of chase (Figure 2B; *Cre*-: 6.60 ± 1.32 *vs. Cre*+: 9.58 ± 3.37% total; *p* = 0.064, Mann Whitney test, *n* = 4-5 animals). The secreted labelled proinsulin:insulin ratio was also increased after 1.5 h of chase (Figure 2C; *Cre*-: 0.85 ± 0.40 *vs. Cre*+: 2.05 ± 0.88; *p*<0.05, Mann Whitney test, *n* = 4-5 animals), but no apparent change was noted in the labelled proinsulin:insulin ratio inside the cells (Figure 2D; 1.5 h: *Cre*-: 0.29 ± 0.07 *vs. Cre*+: 0.27 ± 0.04, 4 h: *Cre*- : 0.17 ± 0.06 *vs. Cre+* : 0.16 ± 0.04, ns, Mann Whitney test, *n* = 4-5 animals) or in the newly synthesised stored insulin remaining in the cells (Figure 2E; 1.5 h: *Cre*-: 0.56 ± 0.09 *vs. Cre*+: 0.61 ± 0.07; 4 h: *Cre*- : 0.45 ± 0.15 *vs. Cre+* : 0.53 ± 0.07, ns, Mann Whitney test, *n* = 4-5 animals).

**Figure 2:**
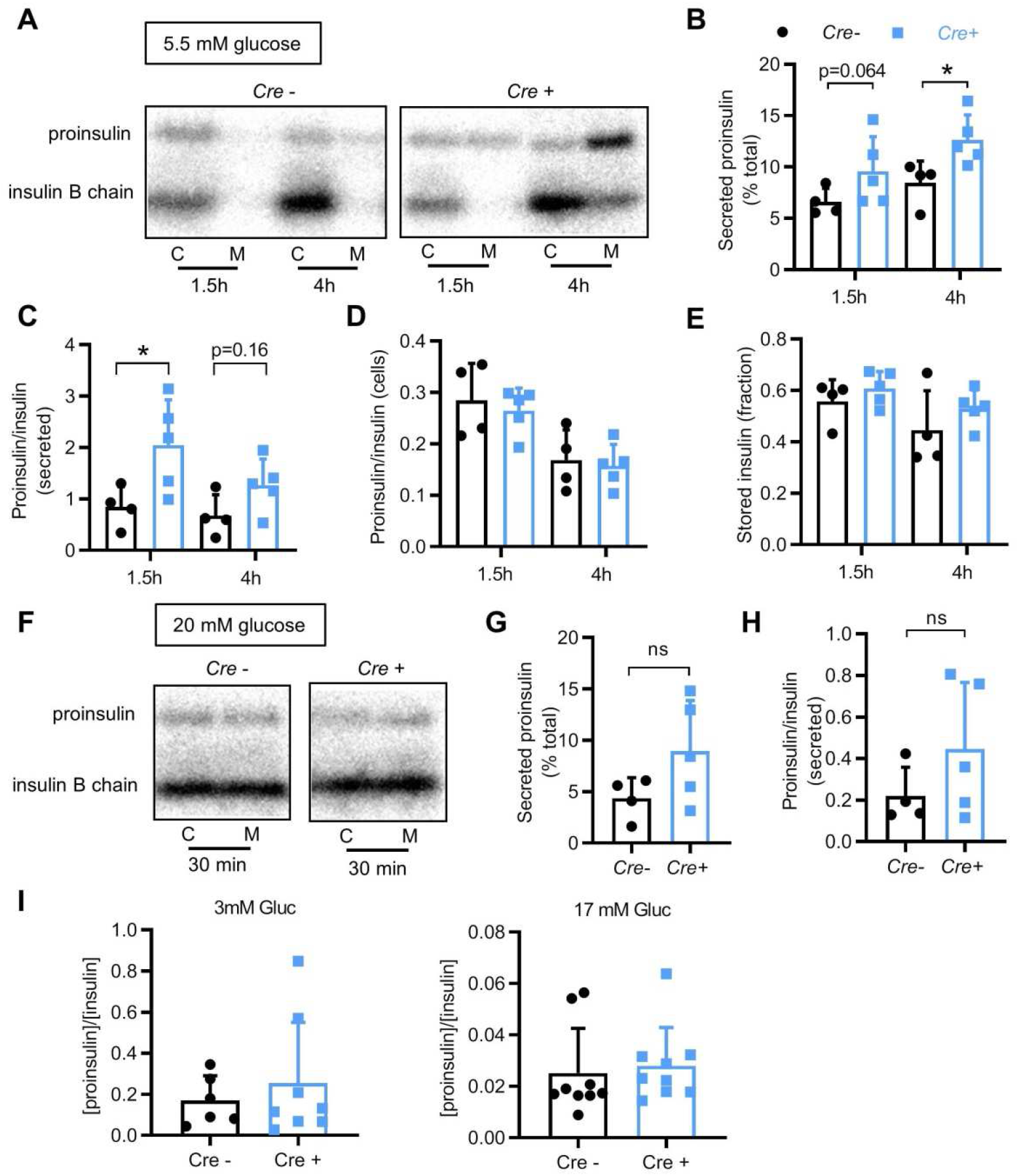
Deletion of Stard10 increased basal secretion of newly synthesised proinsulin but did not affect total secreted proinsulin:insulin ratio. A, Representative phosphorimages from reducing gels representing ^35^S-labelled proinsulin and insulin (B chain) originating from cell lysate (C) or secreting medium (M) samples after 1.5 or 4 h of chase in 5.5 mM glucose medium. B, Secreted proinsulin after 1.5 or 4 h of chase in 5.5 mM glucose medium expressed as percentage of total labelled proinsulin (*n* = 4-5 animals; *p<0.05, Mann Whitney test). C, Secreted proinsulin:insulin ratio after 1.5 or 4 h of chase in 5.5 mM glucose medium (*n* = 4-5 animals; *p<0.05, Mann Whitney test). D, Fraction of stored processed insulin remaining inside the cells after 1.5 or 4 h of chase in 5.5 mM glucose medium (*n* = 4-5 animals; ns, Mann Whitney test). E, Cellular proinsulin:insulin ratio after 1.5 or 4 h of chase in 5.5 mM glucose medium (*n* = 4-5 animals; ns, Mann Whitney test). F, Representative phosphorimages from reducing gels representing ^35^S-labelled proinsulin and insulin (B chain) originating from cell lysate (C) or secreting medium (M) samples after 30 min. of chase in 20 mM glucose medium. G, Secreted proinsulin after 30 min. of chase in 20 mM glucose medium expressed as percentage of total labelled proinsulin (*n* = 4-5 animals; ns, Mann Whitney test). H, Secreted proinsulin:insulin ratio after 30 min. of chase in 20 mM glucose medium (*n* = 4-5 animals; ns, Mann Whitney test). Quantification for B, C, D, E, G and H was done on the phosphorimages obtained from reducing gels. I, Secreted total proinsulin:insulin ratio after 30 min. secretion by isolated islets in 3 or 17 mM glucose Krebs-HEPES-bicarbonate buffer.

A similar experiment was carried out where the islets were chased after 30 min. in 20 mM glucose (Figure 2F). Although the proinsulin secretion and the secreted proinsulin:insulin ratio from the KO islets at 20 mM glucose tended to be increased *vs* wild-type islets, values were not significantly different between genotypes (Figure 2G and H). It is worth noting that insulin granules ordinarily have a long lifespan in β-cells, with a half-life of several days, indicating that the labelled peptides observed in this experiment originate from young granules.

In order to determine whether the increase in newly synthesised proinsulin secretion observed in the pulse-chase experiment reflected a change in total secreted proinsulin, we measured total insulin and proinsulin in the secretion medium after 30 min. in low (3 mM) or stimulating (17 mM) glucose concentrations. However, the secreted proinsulin:insulin ratios remained unchanged between genotypes at both glucose concentrations (Figure 2I).

### 3.4. Preserved glucose-regulated membrane potential and β-cell-β-cell connectivity in Stard10KO islets

Our previous study [10] showed that deletion of *Stard10* in the β-cell impaired glucose-induced cytoplasmic Ca^2+^ increases and insulin secretion. To test for a potential upstream defect, e.g. in the closure of ATP-sensitive K^+^ (K_ATP_) channels [24], we performed perforated patch-clamp electrophysiology [25] to measure plasma membrane potential in dispersed single β-cells. No change in glucose-induced membrane depolarisation was observed in β-cells from β*Stard10*KO animals compared to WT littermates (Supplemental Figure 2A,B).

β-cell-β-cell connections are essential for synchronised intra-islet Ca^2+^ influx, and ultimately efficient insulin secretion [4]. We subjected the individual Ca^2+^ traces recorded from fluo-2-loaded β-cells in the intact mouse islets to correlation (Pearson R) analysis to map cell-cell connectivity [7,8,26]. In the presence of low (3 mM) glucose, β-cells displayed low levels of coordinated activity in islets of WT and whole body *Stard10*KO animals, as assessed by counting the numbers of coordinated cell pairs (Supplemental Figure 3A,B; 21.70 ± 9.13% *versus* 18.95 ± 17.47% for WT *versus* KO, respectively, ns). By contrast, β-cells displayed highly coordinated Ca^2+^ responses upon addition of 17 mM glucose or 20 mM KCl (the latter provoking depolarisation and a synchronized Ca^2+^ peak; not shown) in WT islets. None of the above parameters were altered in KO islets (Supplemental Figure 3A,B; 17 mM G: 98.61 ± 1.34% *versus* 92.23 ± 10.46% for WT *vs* KO; KCl: 92.63 ± 7.23% *vs* 93.36 ± 11.44% for WT *vs* KO, respectively; ns). Similarly, analysis of correlation strength in the same islets (Supplemental Figure 3C,D) revealed no significant differences between genotypes.

### 3.5. Altered lipidomic profile in βStard10KO islets

To determine whether the observed changes in secretory granule ultrastructure (Figure 1A) might result from an altered lipid composition in *Stard10*-null β-cells, we performed targeted mass spectrometry. Total cholesteryl esters (*Cre*-: 1.60 ± 0.36 *vs. Cre*+: 2.24 ± 0.50% total lipids; \*\**p*<0.01, paired t-test) and phosphatidylinositols (PI) (*Cre*-: 7.63 ± 1.15 *vs. Cre*+: 8.98 ± 1.16% total lipids; \**p*<0.05, paired t-test) as well as a particular species of phosphatidylethanolamine (PE 34:0; *Cre*-: 0.019 ± 0.0029 *vs. Cre*+: 0.024 ± 0.0022% total lipids; \**p*<0.05, paired t-test) were all significantly increased in the KO islets vs. WT (Figure 3), suggesting that STARD10 is involved in the regulation of the turnover of these lipid species.

**Figure 3:**
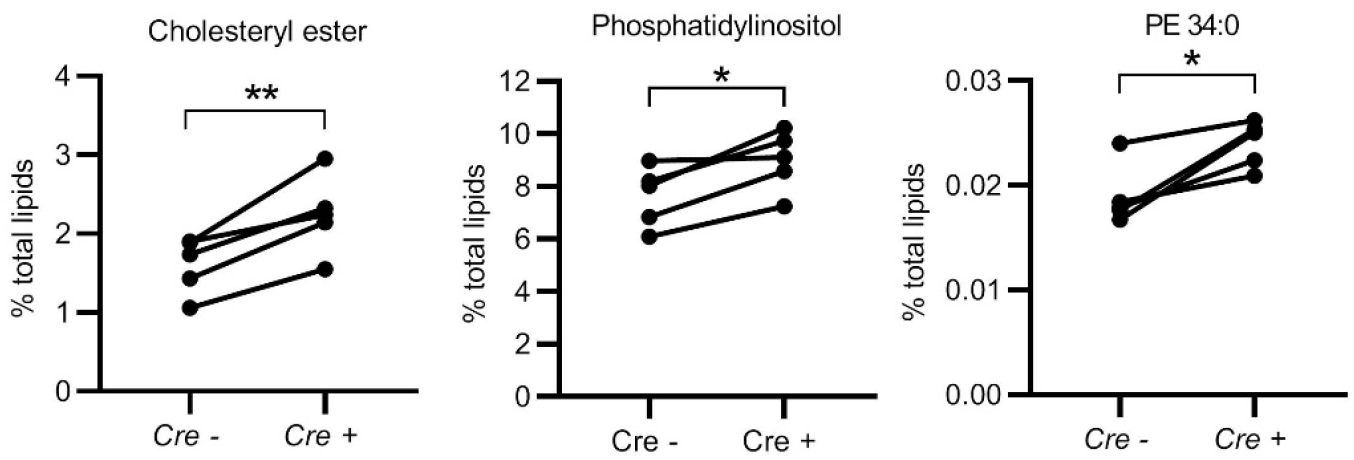
Altered lipidomic profile in βStard10KO islets. The deletion of *Stard10* in mouse pancreatic β-cells significantly increased total cholesteryl esters, phosphatidylinositols and the phosphatidylethanolamine species 34:0 in islets (n = 5 animals; *p<0.05, **p<0.01, paired *t*-test)

### 3.6. Altered expression of genes controlling Ca^2+^ fluxes, exocytosis and inositol phospholipid signalling in βStard10KO islets

In order to assess the effect of β-cell selective *Stard10* deletion on the transcriptome, we performed massive parallel RNA sequencing (RNAseq) on islets from WT and β*Stard10*KO mice. We identified 88 differentially regulated genes (*padj*<0.05, *n* = 6 animals), with 33 upregulated and 55 dowregulated in KO *versus* WT islets. The list of the 20 most significantly regulated genes is presented in Figure 4A and Table 1 (*padj* ranging from 1.71×10^−157^ to 2.25×10^−3^). As expected, a decrease in *Ins1* expression in the *Cre+* mice carrying an Ins1-*Cre* knock-in allele was observed. Interestingly, *Scg2*, encoding secretogranin 2, a member of the granin protein family, localised in secretory vesicles [27], and *Rasd2*, also known as Ras homologue enriched in striatum (Rhes), were upregulated. Suggesting possible changes in phosphoinositide signalling, *Pirt* (phosphoinositide-interacting regulator of transient receptor potential channels), a regulator of transient receptor potential Ca^2+^ channels [28] and *Syt1* (synaptotagmin 1) regulating fast exocytosis and endocytosis in INS1 cells [29] were downregulated in β*Stard10*KO islets.

**Figure 4:**
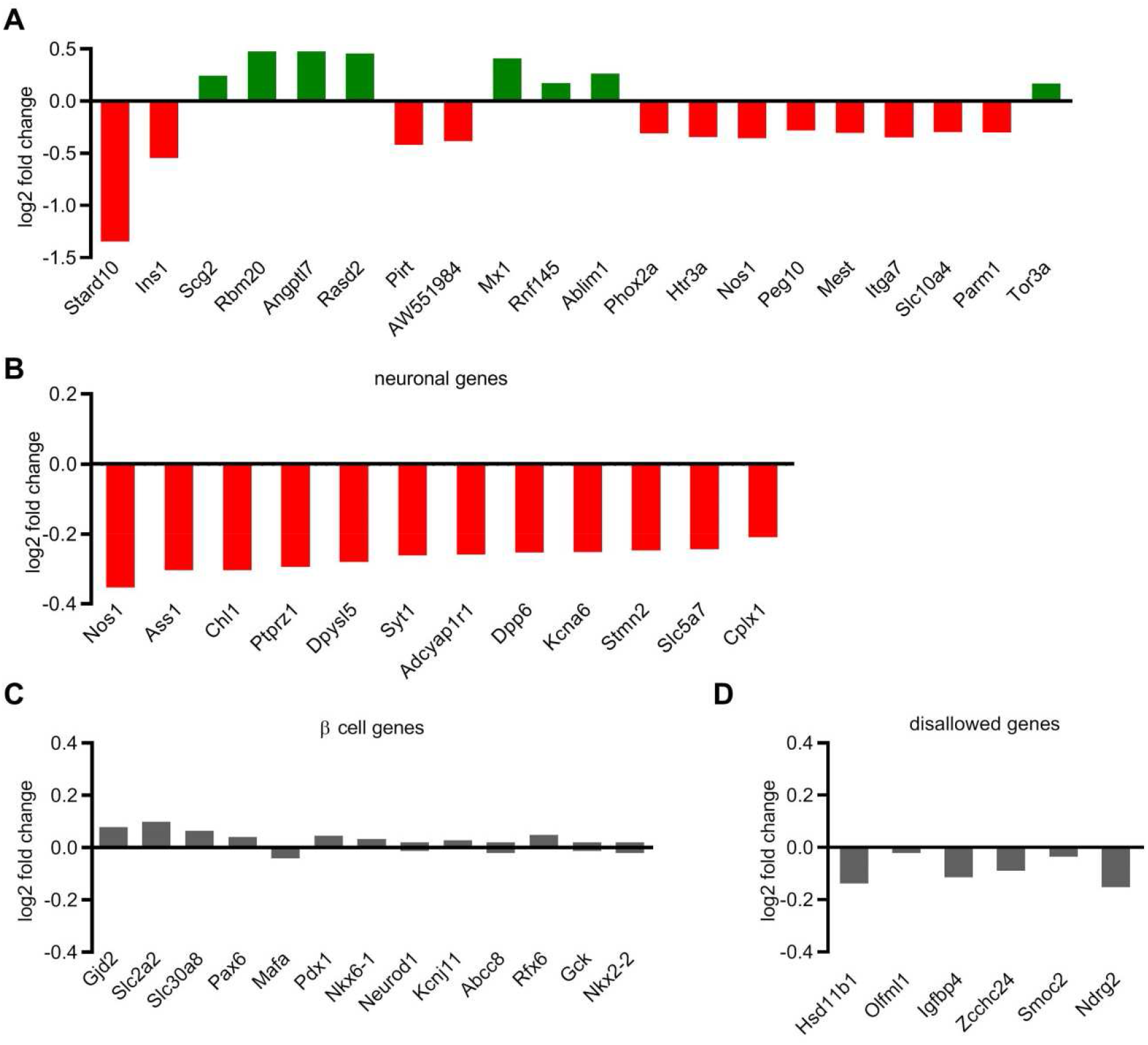
*Identification of differentially-regulated genes in βStard10KO islets* by *RNA seq*. A, 20 first differentially regulated genes in *βStard10KO*, ranked by increasing adjusted p-value with their relative expression *versus* WT (log2 fold change, n = 6 animals per genotype) B, Genes identified by the GO consortium as component of “neuron projection” and “neuron part”, found enriched among the downregulated genes identified in the *βStard10KO* islets, shown as relative expression *versus* WT (log2 fold change). Neither β-cell signature genes (C) nor “disallowed” β-cell gene family (D) were differentially regulated in *βStard10KO vs*. WT islets.

**Table 1.** RNA seq identification of differentially regulated genes in βStard10KO islets. 20 first differentially regulated genes in *βStard10KO*, ranked by increasing adjusted p-value (padj) with their relative expression *versus* WT (log2 fold change)

|  | symbol | log2FoldChange | pvalue | padj | entrezID |
| --- | --- | --- | --- | --- | --- |
| ENSMUSG00000030688 | Stard10 | -1.3455 | 1.13E-161 | 1.71E-157 | 56018 |
| ENSMUSG00000035804 | Ins1 | -0.5451 | 1.46E-17 | 1.11E-13 | 16333 |
| ENSMUSG00000050711 | Scg2 | 0.2439 | 2.13E-13 | 1.07E-09 | 20254 |
| ENSMUSG00000043639 | Rbm20 | 0.4753 | 3.35E-12 | 1.27E-08 | 73713 |
| ENSMUSG00000028989 | Angptl7 | 0.4761 | 1.40E-10 | 4.23E-07 | 654812 |
| ENSMUSG00000034472 | Rasd2 | 0.4568 | 2.99E-10 | 7.52E-07 | 75141 |
| ENSMUSG00000048070 | Pirt | -0.4203 | 1.71E-09 | 3.68E-06 | 193003 |
| ENSMUSG00000038112 | AW551984 | -0.3837 | 5.31E-09 | 1.00E-05 | 244810 |
| ENSMUSG00000000386 | Mx1 | 0.4094 | 3.87E-08 | 6.49E-05 | 17857 |
| ENSMUSG00000019189 | Rnf145 | 0.1714 | 1.22E-07 | 0.000185 | 74315 |
| ENSMUSG00000025085 | Ablim1 | 0.2650 | 1.69E-07 | 0.000232 | 226251 |
| ENSMUSG00000007946 | Phox2a | -0.3054 | 2.01E-07 | 0.000253 | 11859 |
| ENSMUSG00000032269 | Htr3a | -0.3423 | 2.37E-07 | 0.000276 | 15561 |
| ENSMUSG00000029361 | Nos1 | -0.3533 | 4.34E-07 | 0.000468 | 18125 |
| ENSMUSG00000092035 | Peg10 | -0.2823 | 8.04E-07 | 0.00081 | 170676 |
| ENSMUSG00000051855 | Mest | -0.3041 | 1.06E-06 | 0.000999 | 17294 |
| ENSMUSG00000025348 | Itga7 | -0.3500 | 1.38E-06 | 0.001223 | 16404 |
| ENSMUSG00000029219 | Slc10a4 | -0.2939 | 1.53E-06 | 0.00128 | 231290 |
| ENSMUSG00000034981 | Parm1 | -0.3002 | 1.89E-06 | 0.001502 | 231440 |
| ENSMUSG00000060519 | Tor3a | 0.1692 | 2.98E-06 | 0.002254 | 30935 |

Enrichment analysis through the gene ontology consortium website (http://www.geneontology.org/) identified genes whose products are localised in neuronal projections (11 genes, fold enrichment = 4.57, *p*<0.05) and neurons (12 genes, fold enrichment = 3.95, *p*<0.05) among the down-regulated genes (Figure 4B, Supplemental Table 2). As secretory cells, neurons share their transport and secretory machinery with β-cells, suggesting that these genes play a role in the insulin secretion process. Among the genes identified, several are implicated in insulin secretion and/or β-cell survival: *Chl1* (cell adhesion molecule L1-like), whose silencing has previously been shown to reduce glucose-induced insulin secretion in INS1 cells [30] and is down regulated in islets from Type 2 diabetic subjects [31]; *Nos1* (neuronal nitric oxyde synthase), implicated in insulin secretion and β-cell survival [32]; *Adcyap1r1* (adenylate cyclase activating polypeptide 1 receptor 1), encoding the PAC1 receptor of the pituitary adenylate cyclase activating polypeptide (PACAP) neuroendocrine factor [33] and *Cplx1* (complexin 1) [34]. None of several key β-cell signature genes involved in the regulation of *Ins* expression and β-cell identity (Figure 4C), or of the “disallowed” β-cell gene family (Figure 4D) [3], were affected by *Stard10* deletion.

### 3.6. STARD10 interactome analysis identifies proteins involved in phosphatidylinositide signalling

Protein interacting partners of STARD10 were identified by immunoprecipitation of the endogenous protein in the rat pancreatic β-cell line INS1 (832/13) and liquid chromatography–tandem mass spectrometry analysis after tandem mass tag (TMT) labelling of the digested samples. 303 significantly enriched proteins were detected in the STARD10 pull-down condition compared with the non-targetting control IgG (*p*<0.05 and FDR<0.05, paired t-test) (Table 2). Among these significant interactions, we observed proteins well known to play a role in β-cell physiology: the pore forming subunit Kir6.2 (Kcnj11, ATP-sensitive inward rectifier potassium channel 11) and its auxiliary subunit, the sulfonylurea receptor SUR1 (Abcc8, ATP-binding cassette sub-family C member 8) of the ATP-sensitive K^+^ (K_ATP_) channel [35]; the phosphoinositide PI(4,5)P_2_-interacting protein Granuphilin (Sytl4; synaptotagmin-like 4) associates with and is involved in the docking of the insulin granules [36]; the polypyrimidine tract-binding protein 1 (Ptbp1) is required for the translation of proteins localised in the insulin granule in response to glucose [37]. Gene ontology analysis of the identified STARD10 binding partners reflected an enrichment in proteins associated with RNA binding and processing, gene expression and splicing (Supplemental Tables 3, 4 and 5).

**Table 2.**
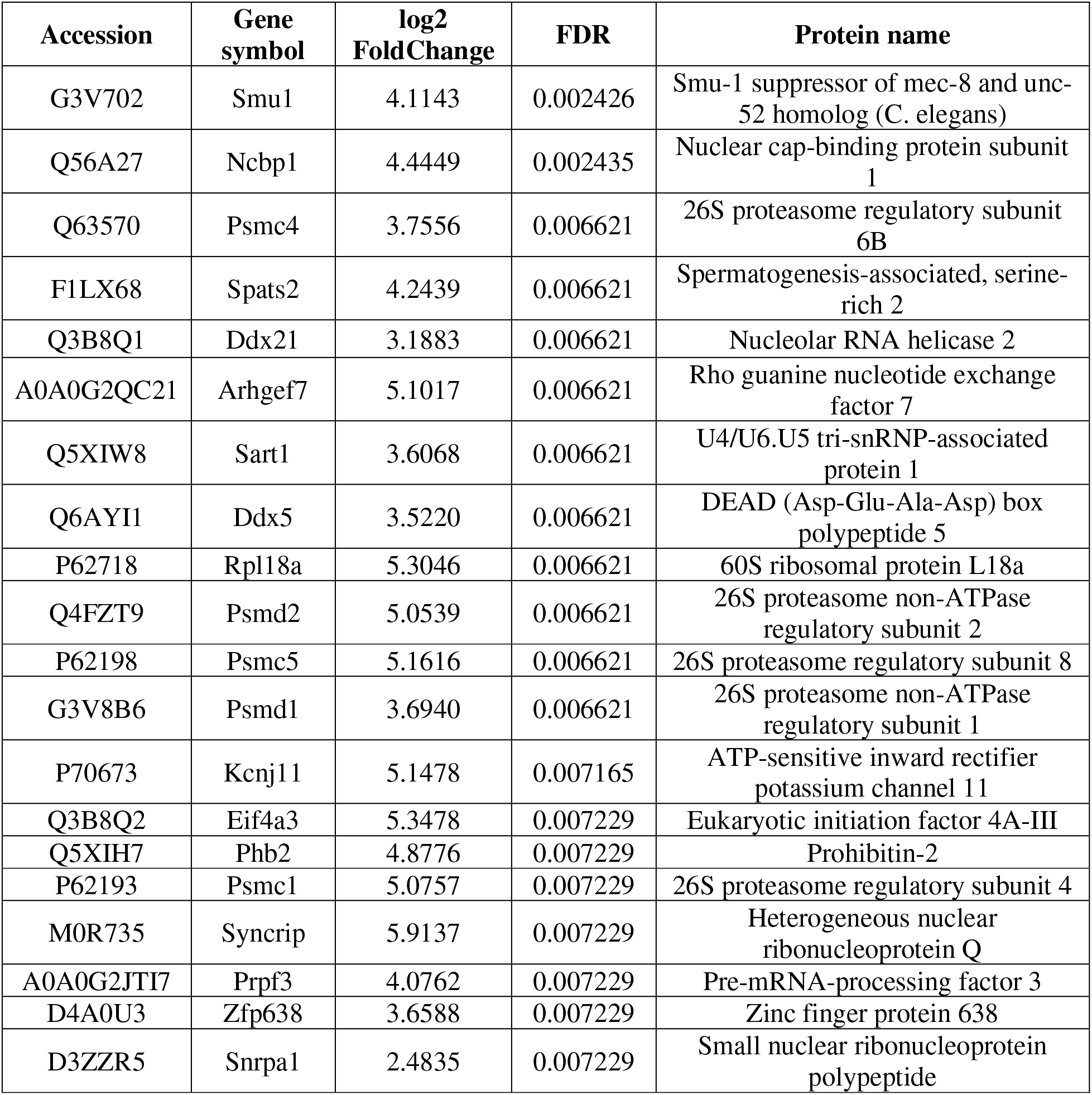
STARD10 binding partners identified by mass spectrometry in INS1 (832/13) cells. 20 most significantly enriched proteins found in the STARD10 pull down condition versus control IgG with a cut-off of at least 5 unique peptides identified.

Interestingly, STARD10 also interacted with the inositol lipid kinase phosphatidylinositol 5-phosphate 4-kinase type-2 gamma (Pip4k2c) enzyme, which converts phosphatidylinositol 5-phosphate (PI(5)P) into phosphatidylinositol 4,5-bisphosphate (PI(4,5)P_2_) [38].

### 3.7. STARD10 binds to phosphatidylinositides

To assess which membrane lipids may be bound by STARD10, we generated and purified the recombinant protein from a bacterial expression construct comprising the human STARD10 coding sequence fused with a 6His-MBP (Maltose Binding Protein) tag. Following recombinant protein production in *E. coli* and protein purification, we performed a lipid overlay assay, in which *in vitro* lipid binding to a range of phospholipids spotted onto a nitrocellulose membrane is assessed [39]. The efficiency and specificity of protein purification was monitored by SDS-PAGE staining with Coomassie blue (Figure 5A). Purified STARD10 protein was incubated with the PIP membrane strips and detected using an anti-STARD10 antibody and secondary anti-goat-HRP antibody (Figure 5B). Interestingly, STARD10 interacted with all PIP species interrogated, suggesting that STARD10 may bind to membranes containing these phospholipids.

**Figure 5.**
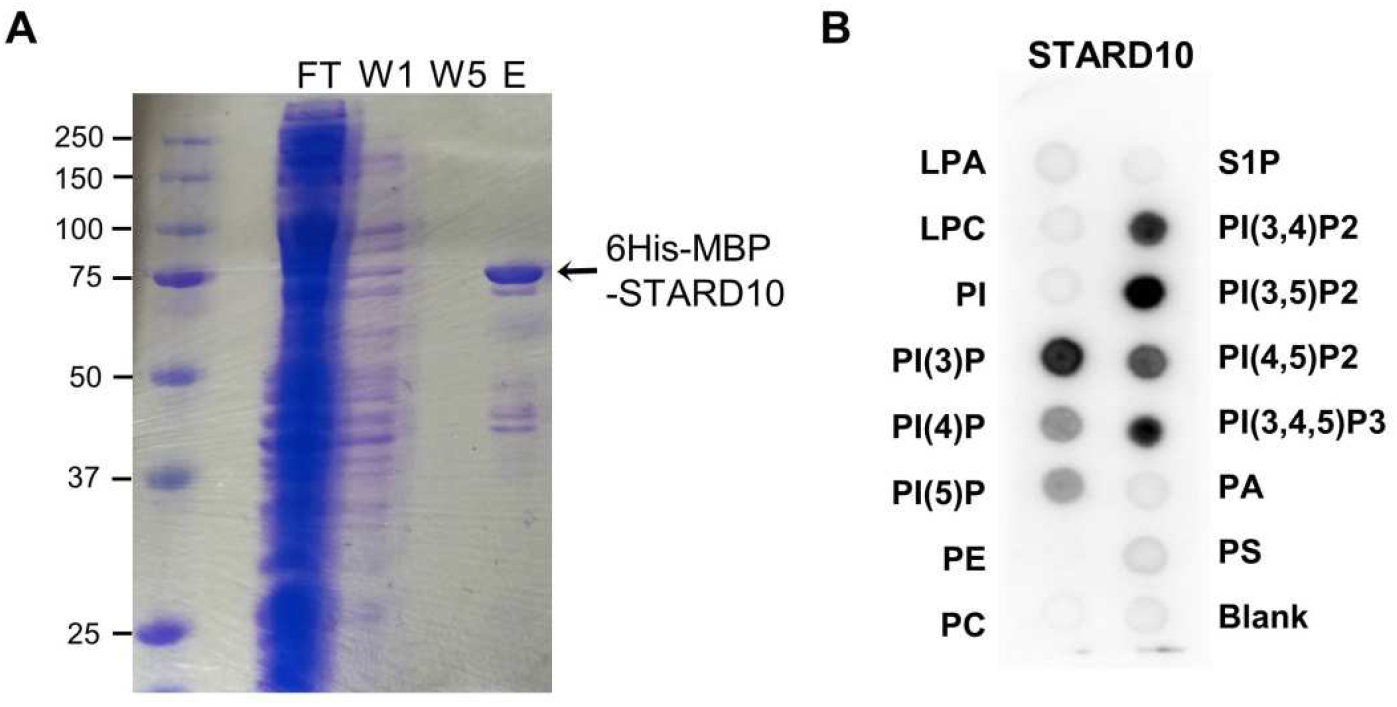
Phosphoinositide binding to STARD10. A, Coomassie blue staining of 6His-MBP-STARD10 (75kDa) purified by Immobilised Metal Affinity column (IMAC): FT: Flow-Through, W1: first column wash, W5: last column wash, E: elution. B, Lipid overlay assay. LPA, lysophosphatidic acid; S1P, Sphingosine-1-phosphate; LPC, Lysophosphatidylcholine; PI, Phosphatidylinositol; PI(3)P, PI-(3)-phosphate; PI(4)P, PI-(4)-phosphate; PI(5)P, PI-(5)-phosphate; PE, Phosphatidylethanolamine; PC, Phosphatidylcholine; PI(3,4)P_2_, PI-(3,4)-bisphosphate; PI(3,5)P_2_, PI-(3,5)-bisphosphate; PI(4,5)P_2_, PI-(4,5)-bisphosphate; PIP(3,4,5)P_3_, PI-(3,4,5)-trisphosphate; PA, Phosphatidic acid; PS, Phosphatidylserine. Immunodetection of bound protein was performed using a primary anti-STARD10 antibody (Santa Cruz) and a secondary anti-goat-HRP antibody (Santa Cruz). STARD10 bound to all PIP species.

### 3.8. Resolution of STARD10 structure and molecular docking identify PI and PI(3)P as potential ligands

The above lipid overlay assay reveals interaction with phospholipids but does not distinguish between surface binding and uptake of the lipid into the lipid binding pocket of the protein. In order to explore the potential binding of inositol phospholipids in this pocket, we sought to obtain a crystal structure of the purified protein. Purified STARD10 generated a well diffracting crystal. The crystal structure was then resolved at 2.3 Å resolution by molecular replacement using the structure of STARD5 (PDB ID: 2R55) (Figure 6A; Supplemental movie 1; Supplemental Table 6).

In the STARD10 crystal structure, the dimensions of the cavity are such that the protein is expected to readily accommodate phosphatidylinositols, whereas STARD2 cavity, due to its smaller size, gives poor predicted binding energy. On this basis, STARD10 binding pocket is expected to bind PI and various phosphorylated PIs with higher affinity than STARD2.

**Figure 6.**
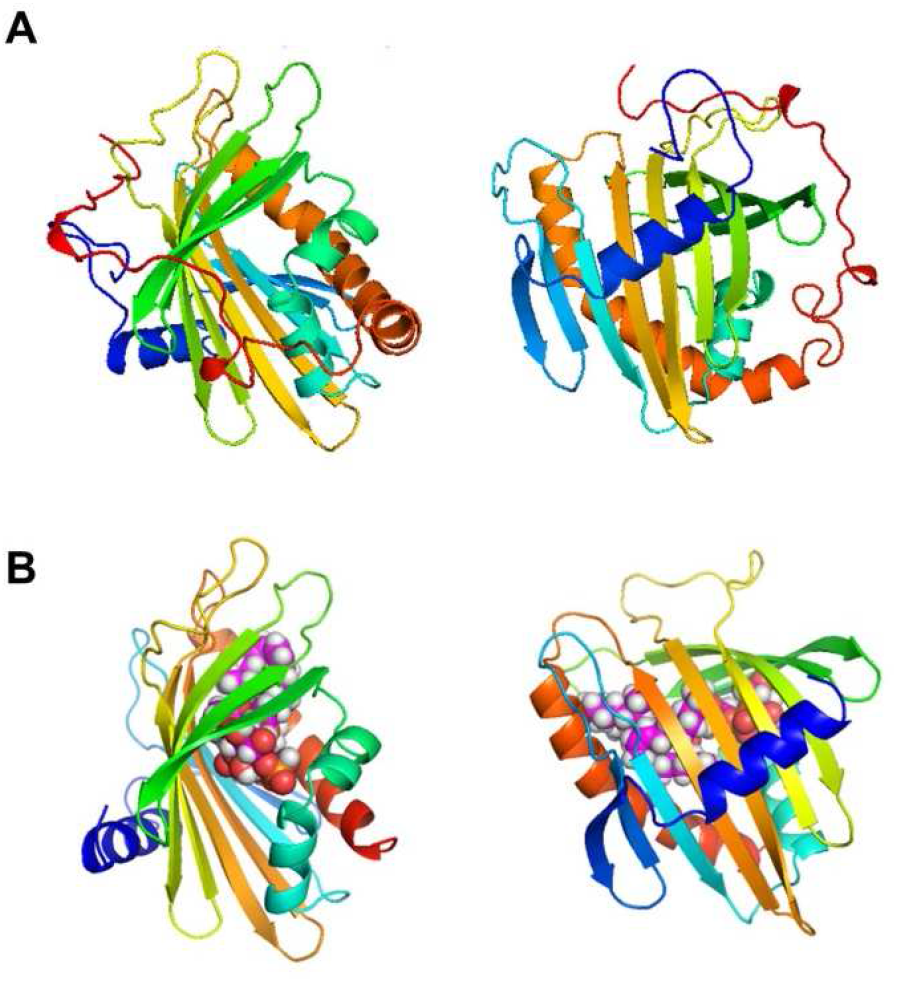
Structure of H. sapiens STARD10. A, Three-dimensional view of the crystal structure of unliganded human STARD10: ribbon diagram colored from the N-terminus (blue) to the C-terminus (red). B, Docking of phosphatidyl-inositol 3 phosphate (PI(3)P) to the human STARD10 structure. STARD10 cavity is larger than the phosphatidylcholine transporter protein STARD2 and contrary to the latter readily accommodates phosphatidylinositols.

The head groups of PI and PI(3)P (glycero-inositol-1-phosphate and glycero-3-phosphoinositol-1-phosphate) were then docked into the 3-dimensional structure of STARD10 (described above) and the closer family member STARD2 [40] by using the Bristol University Docking Engine (BUDE) [41] (Figure 5B, supplemental movie 2). The software was used to dock different representative conformations of the ligand to the target protein: 87 conformers for PI and 85 for PI(3)P. When these conformers were docked to STARD10, all final poses were located inside the binding cavity. On the other hand, for STARD2, only two conformers were placed by BUDE inside the protein, the rest were docked onto the outer surface of the protein, suggesting that, contrary to STARD10, binding inside STARD2 by PI and PI(3)P is unfavourable. Likewise, the predicted binding energies of PI and PI(3)P headgroups to STARD10 is about 40 kJ mol^−1^ better than those to STARD2.

To address which intracellular membranes may be affected by alterations in STARD10 expression, and hence lipid distribution, we next performed confocal analysis of the subcellular localization of STARD10 tagged at the carboxy- or amino-terminus with green fluorescent protein (STARD10-GFP or GFP-STARD10). However, this indicated a largely homogeneous cytosolic and nuclear distribution (Supplemental Figure 4), precluding the identification of likely target membranes or organelles.

## 4. Discussion

Our earlier demonstration [10] that *STARD10* is the likely causal gene at the T2D risk locus on chromosome 11q13 has emphasised the probable importance of lipid transfer for the normal physiology of the pancreatic β-cell. Indeed, the role of lipid transfer proteins has emerged recently as an important element in cell biology [42].

Several lines of evidence presented in our studies now support a role for STARD10 in PI/PIPs binding and potentially transport between intracellular membranes: (a) Resolution of the crystal structure of STARD10 and molecular docking predict binding of PI/PIPs in its binding pocket (Figure 6). (b) Purified recombinant STARD10 binds to phosphatidylinositide species in a lipid overlay assay (Figure 5). (c) The phosphatidylinositol content was increased in βStard10KO compared to control islets (Figure 3). (d) The expression of the phosphoinositide binding proteins *Syt1* and *Pirt* were decreased in βStard10KO islets (Figure 4). *Pirt* encodes a phosphoinositide interacting regulator of the transient receptor potential (TRP) Ca^2+^ channels. Several TRP channels are expressed in the β-cell and an increasing amount of data points towards a role of these channels in the regulation of insulin secretion [43]. A previous study [44] has shown that female *Pirt*^-/-^ mice are heavier than controls and develop glucose intolerance. (e) Our proteomic study identified the phosphatidylinositol 5-phosphate 4-kinase type-2 gamma (Pip4k2c) enzyme and the PI(4,5)P_2_-binding protein granuphilin [45] as STARD10 binding partners.

Taken together, these data suggest that STARD10 may bind PI and/or PIPs either in the lipid binding pocket, as predicted by the crystal structure and molecular docking (Figure 6), or at the surface of the protein. We note that the lipid overlay assay is consistent with a binding of STARD10 to membranes containing PIP lipids (with an affinity which appears higher for (3)P containing species). This could be important for the targeting of STARD10 to specific membrane compartments. In contrast to its closest relative STARD2, which has been shown to selectively bind phosphatidylcholine [46], our crystal structure determination and molecular docking indicate that STARD10, due to its larger binding pocket, is predicted to bind phosphatidylinositol (PI) and its various phosphorylated forms (PIPs) (Figure 6) in addition to its previously recognised ligands phosphatidylcholine and phosphatidylethanolamine [12].

Consistent with likely mis-targeting of lipids to membranes involved in insulin secretion, we demonstrate that STARD10 is required for normal secretory granule maturation in murine β-cells. Of note, recent findings [47] have indicated that cholesterol is also required for normal granule maturation. In addition, in INS-1 (832/13) cells, the insulin secretory granule was found to contain a high proportion of phosphatidylinositol (~20%, 5-fold that in the whole cell) [48]. In contrast, we were not able to obtain evidence for an effect of STARD10 variation on human β-cell granules, perhaps reflecting a greater diversity in granule structure observed in man, as well as the limited number of subjects available for analysis.

RNAseq analysis of the transcriptome of WT and *βStard10*KO islets identified differentially-expressed genes which are likely to play a role in adult β-cell physiology (Figure 4). These include *Syt1* (synaptotagmin 1) and *Cplx1* (complexin 1), whose genes products are part of the molecular machinery driving vesicle exocytosis and form a tripartite complex with Soluble N-ethylmaleimide sensitive factor Attachment protein Receptor (SNARE) proteins [49]. However, none of these genes are readily linked to a change in insulin crystallisation or granule biogenesis. Importantly, we also identified *Pirt*, a phosphoinositide-interacting regulator of transient receptor potential channels, as a binding partner of STARD10 and thus likely either to sense or to influence phosphoinositides levels in cells.

Proteomic analysis of the STARD10 binding partners in INS1(832/13) pancreatic β-cells (Table 2) identified several proteins involved in insulin secretion such as the subunits of the K_ATP_ channel Kir6.2 and SUR1 and the secretory vesicle protein granuphilin, which possesses a C2 domain capable of binding the phosphoinositide PIP_2_ [50]. Another binding partner, PTBP1 (Polypyrimidine tract-binding protein 1), regulates the translation of insulin granule proteins in response to glucose [37]. The potential regulation by STARD10 of the activity of this protein could conceivably affect granule structure.

We also considered the possibility that altered granule composition may reflect altered intracellular Zn^2+^ levels given that similar changes are observed after deletion of the gene encoding the Zn^2+^ transporter ZnT8, *Slc30a8* [51]. Thus, total Zn^2+^ content, likely reflecting intragranular Zn^2+^, assessed by ICP/MS, was increased in islets of β*Stard10*KO *vs*. WT mice (Figure 1). The latter appears unlikely to be due to changes in the expression of *ZnT8* (encoded by *Slc30a8*), which we did not observe, but might reflect altered intrinsic activity, or granule recruitment, of ZnT8 due to altered membrane lipid composition [52]. Consistent with this possibility, anionic phosphatidylinositols increase, whereas non bilayer phospholipids (promoting membrane curvature) lysophosphatidylcholine (LPC) and phosphatidylethanolamine (PE), lower ZnT8 activity [52].

An unexpected finding from the current study is that β*Stard10*KO islets secreted more radiolabeled proinsulin versus WT after 4 h of chase in 5.5 mM glucose medium, whilst no difference was observed between genotypes at 20 mM glucose. These results contrast with a *decrease* in the proinsulin:insulin ratio observed in plasma from β*Stard10*KO mice [10] and a similarly lowered ratio in human risk variant carriers versus controls [22]. One possible explanation is that increased proinsulin secretion in unstimulated cells may enrich for storage of mature, processed insulin, hence producing a lowered proinsulin:insulin release during stimulation with high glucose. According to studies using radioactive labelling [53] or live cell imaging of a SNAP-tag fused with insulin [54], young, newly synthesised insulin granules are preferentially secreted upon glucose stimulation. Given their reduced transit time, the proportion of unprocessed proinsulin in the young granules could be higher than in older ones. Other evidence points towards the existence of vesicular trafficking pathways leading to unstimulated, “constitutive-like” secretion of proinsulin [55]. It has been speculated [56] that this involves routing to the endosomal compartment and either cargo degradation in the lysosome, or secretion at the plasma membrane through a “constitutive-like” pathway [57]. By changing the lipid composition of granules, STARD10 may modulate their trafficking properties, affect the age of secreted granules and/or decrease their targeting to the lysosomes.

A striking finding of our studies is the change in lipid composition observed in the β*Stard10*KO islets (Figure 3). Although it was not possible to assess which intracellular membranes may be affected given the limited sensitivity of the bulk analysis performed, we note that an excess of cholesterol has previously been shown to inhibit insulin exocytosis [58,59] and former studies points towards a role for phosphoinositide signalling in insulin secretion [60,61]. The PI transporter TMEM24 regulates pulsatile insulin secretion by replenishing the phosphoinositide pool at the endoplasmic reticulum-plasma membrane contact sites [62]. It is thus tempting to speculate that STARD10 might affect insulin secretion in a similar fashion.

In conclusion, we identify phosphatidylinositols as potential new ligands for STARD10. Changes in *STARD10* expression in carriers of type 2 diabetes risk alleles may consequently affect the lipid composition, alter β-cell granule maturation, and, ultimately, insulin synthesis and secretion.

## Author contributions

G.R.C. designed and conducted the in vitro studies and contributed to the writing of the manuscript. E.H. performed the electrophysiology studies. L.H. and P.A. performed the pulse chase studies and contributed to the discussion. A.T., A.F. A.K. A.M. and M.S. contributed to the electron microscopy studies. A.P. performed the immunocytochemistry studies. K.C., M.H., and D.B.W. produced STARD10 recombinant protein and performed the structural studies. T.J.P. performed the analyses of the RNAseq data. E.G., T.S., D.J.H., L.J.B. V.S and W.D. contributed to the connectivity analysis. N.S.A., P.I.B., F.T. and A.K.K.T. performed the lipidomic analysis. K.J.H and P.A.L. performed the interactome analysis. R.B.S. performed the STARD10 ligand docking studies. F.A. provided the STARD10-GFP and GFP-STARD10 constructs. A.C. performed the ICP/MS analysis I.L. was the holder of the Home Office project licence and responsible for the work carried on animals in this study. G.A.R. conceived the study and co-wrote the manuscript. G.A.R. is the guarantor of this work and, as such, had full access to all the data in the study and takes responsibility for the integrity of the data and the accuracy of the data analysis.

## Acknowledgments

The authors are grateful to Thomas Di Mattia and Laetitia Voilquin from the Institut de Génétique et de Biologie Moléculaire et Cellulaire (IGBMC), France for construction of the STARD10-GFP and GFP-STARD10 plasmids. We thank Rhodri M. L. Morgan for assistance with data collection for the determination of STARD10 structure and we acknowledge the Diamond Light Source for access to beamlines I03. We would like to thank the Electron Microscopy Facility of the MPI-CBG for their support. This work was supported by the Electron Microscopy and Histology Facility, a Core Facility of the CMCB Technology Platform at TU Dresden

## Study approval

All in vivo procedures were approved by the UK Home Office according to Animals (Scientific Procedures) Act 1986 (HO Licence PPL 70/7349) and were performed at the Central Biomedical Service, Imperial College, London, UK.

Pancreatectomised patient samples from the IMIDIA consortium (www.imidia.org; Solimena, M, personnal communication) with appropriate permissions from donors and/or families and approval by the local ethics committee. A written informed consent was received from participants prior to inclusion in the study.

## Data and Resource Availability

The RNAseq raw sequence data on βStard10KO and WT mice will be made available via deposition to ArrayExpress.

Lipidomic mass spectrometry raw data can be provided upon request and will be depositide in MetaboLights.

The resources generated or analyzed during the current study are available from the corresponding author upon reasonable request.

## Abbreviations

BUDE: Bristol University Docking Engine
GFP: Green Fluorescent Protein
GWAS: Genome Wide Association Study
K_ATP_: ATP-sensitive K^+^ channels
HRP: Horseradish Peroxidase
Kir6.2: *Kcnj11*: Potassium Inwardly Rectifying Channel Subfamily J Member 11
KOMP: NIH Knock-Out Mouse Project
ICP/MS: Inductively coupled plasma mass spectrometry
IMPC: International Mouse Phenotyping Consortium
LPC: Lysophosphatidylcholine
MBP: Maltose Binding Protein
PC: Phosphatidylcholine
PE: Phosphatidylethanolamine
PI: Phosphatidylinositol
PI(3)P: Phosphatidylinositol 3-phosphate
PI(4,5)P2: PIP_2_, Phosphatidylinositol 4,5-bisphosphate
PI(5)P: Phosphatidylinositol 5-phosphate
PIP: phosphatidylinositol phosphate, phosphatidylinositide
*Pip4k2c*: phosphatidylinositol 5-phosphate 4-kinase type-2 gamma
*Pirt*: phosphoinositide-interacting regulator of transient receptor potential channels
RNAseq: RNA sequencing
*Ptbp1*: Polypyrimidine tract-binding protein 1
*Slc30a8*: ZnT8, Solute Carrier Family 30 Member 8
SNARE: Soluble N-ethylmaleimide sensitive factor Attachment protein Receptor
STARD10: StAR Related Lipid Transfer Domain Containing 10
*Syt1*: Synaptotagmin 1
*Sytl4*: synaptotagmin-like 4, granuphilin
TMT: Tandem Mass Tag
TRP: Transient receptor potential

## Appendix A. Supplementary data

