## Supplemental data for "The type 2 diabetes gene product STARD10 is a phosphoinositide binding protein that controls insulin secretory granule biogenesis"

**Online Supplemental Materials**

### Supplemental Research Design and Methods

#### *Electrophysiology*

Pancreatic islets were dissociated into single β-cells and plated onto glass coverslips, as previously described (1). Electrophysiological recordings were performed in perforated patch-clamp configuration using an EPC9 patch-clamp amplifier controlled by Pulse acquisition software (Heka Elektronik, Pfalz, Germany). β-cells were identified morphologically and by depolarisation of the membrane potential in response to 17 mM glucose. β-cells were constantly perfused at 32°C with normal saline solution (mM): 135 NaCl, 5 KCl, 1 $MgCl_2$, 1 $CaCl_2$, 10 HEPES and 16.7 glucose (pH 7.4). Recording electrodes had resistances of 8-10 MΩ and were filled with a solution comprised of (mM): 140 KCl, 5 $MgCl_2$, 3.8 $CaCl_2$, 10 HEPES, 10 EGTA (pH 7.2) and 20–25 μg/ml amphotericin B (Sigma-Aldrich).

#### *$Ca^{2+}$ imaging and connectivity analysis*

$Ca^{2+}$ (with Fluo-2-AM, Invitrogen) imaging were performed as previously described (2, 3). Briefly, Calcium imaging was performed in Krebs-HEPES-Bicarbonate (KHB) buffer (40 mM NaCl, 3.6 mM KCl, 0.5 mM $NaH_2PO_4$, 0.2 mM $MgSO_4$, 1.5 mM $CaCl_2$, 10 mM HEPES, 25 mM $NaHCO_3$), saturated with 95% $O_2$/5% $CO_2$ and adjusted to pH 7.4. Islets were incubated at 37 °C 95% $O_2$/5% $CO_2$ for 45 min. in 10 μM Fluo-2-AM (Invitrogen). Islets were then transferred in a perfusion chamber, mounted on a Zeiss Axiovert confocal microscope and perfused continuously at 34-36°C. Images were acquired every 2 s with a Hamamatsu ImagEM camera and Volocity software (Perkin-Elmer) was used for data capture. Significantly correlated cell pairs were measured as described previously (4).

Correlation coefficient (R) and heatmaps were generated using an in house Matlab script (available upon request).

***Generation of GFP-STARD10 and STARD10-GFP constructs***

STARD10 open reading frame (NM_006645) was amplified by PCR from a MCF7 (ATCC HTB-22) cDNA library using the following primers: P1: 5'- CCC CAT GGA GAA GCT GGC GGC CTC T-3' and P2: 5'- TCA GGT GAG CGA GGT GTC GTC GTC G-3', and cloned into the pSTBlue-1 Blunt vector. STARD10 cDNA was then amplified by PCR with the primers: P1 and 5'- CGG TGA GCG AGG TGT CGT CGT C -3', or 5'- GAG AAG CTG GCG GCC TCT ACA -3' and P2. The PCR fragments were phosphorylated and cloned into the DraI and EcoRV sites of the pENTR1A gateway entry vector, to generate entry plasmids compatible with C-terminal and N-terminal fusions, respectively. STARD10 was further cloned by gateway recombination into pEGFP-N1 and pEGFP-C2 destination vectors to generate STARD10-EGFP and EGFP-STARD10 expression vectors, respectively.

***STARD10 subcellular localisation.***

Cells from the human β-cell line EndoC-βH1 were seeded on 24 mm diameter coverslips in a 6 well-plate and transfected with the GFP-STARD10 or STARD10-GFP constructs using Lipofectamine2000. 48 h after transfection, the cells were imaged live on a spinning disk confocal microscope (Nikon Eclipse Ti, Crest spinning disk, 60x oil objective; Cairn instruments).

**Supplemental figures**

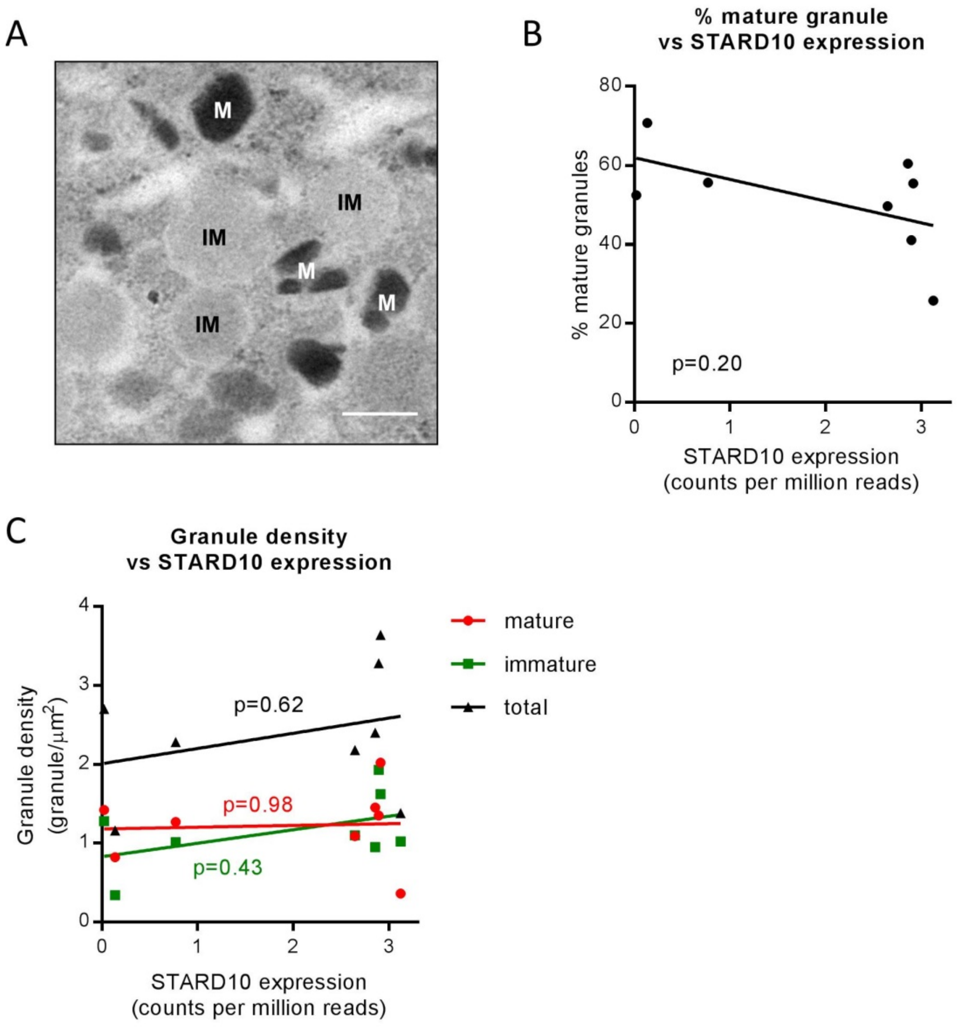


***Supplemental Figure 1. Association of STARD10 expression and granule morphology in human islets***

A, Representative Transmission Electron Microscopy image of a human β-cell taken from a partial pancreatectomy sample showing examples of mature (M) and immature (IM) insulin secretory granules. Scale bar = 200 nm. B, No significant correlation could be observed between the percentage of mature insulin granules and the level of *STARD10* expression measured by RNASeq (n = 8 human donors; ns by non-parametric Spearman correlation, p value shown on graph). C, Identical to B but with density of mature, immature and total insulin granules (granule/μm$^2$) (n = 8 human donors; ns by non-parametric Spearman correlation, p-value shown on graph).

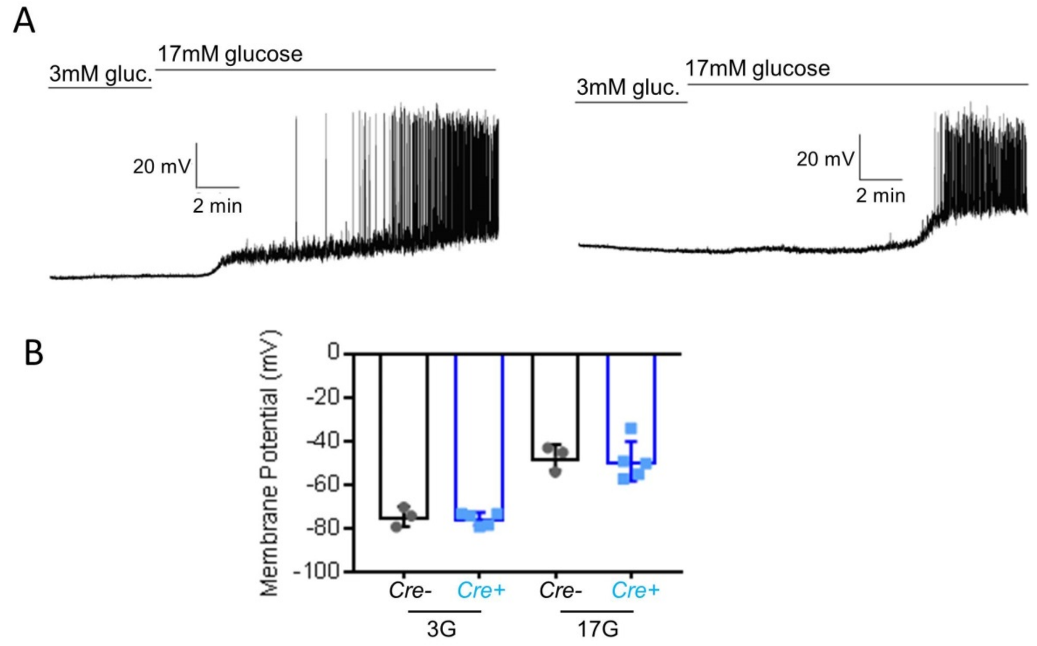


***Supplemental Figure 2*: *Deletion of Stard10 does not affect membrane potential in mouse islet.***

A, Representative current clamp recordings of isolated β cells from wt and β*Stard10*KO mice in response to a 3 and 17 mM glucose application. B, Mean membrane potential at 3 and 17 mM glucose (*n* = 4 to 5 cells from 2 animals per genotype, ns by Student's t-test).

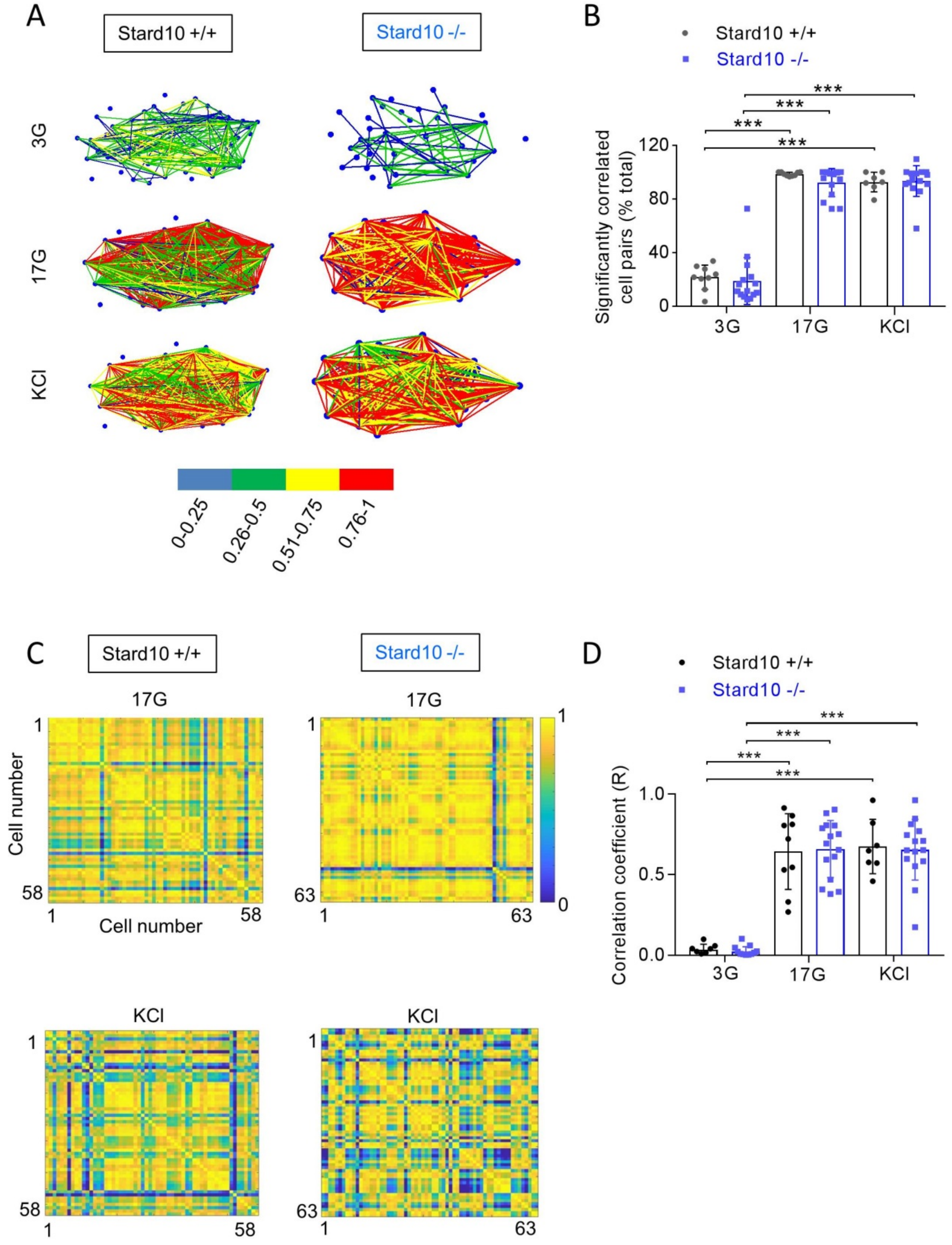


***Supplemental Figure 3: Impact of StarD10 deletion on intercellular connectivity***

A, Representative maps of the recorded cells with coded connectivity strength. The maps show a Cartesian representation of the cells. A colour coded line connects two cells according to the strength of the Pearson R statistic. B, Percentage of correlated cell pairs at 3, 17 mM glucose or 20 mM KCl (*n* = 7 to 15 islets per genotype; ns by 2-way ANOVA). C, Representative heatmaps depicting connectivity strength (Pearson R statistic) of all cell pairs (R values color coded from -1 to 0, blue to yellow respectively). D, The average correlation coefficient (R) between β-cells rises significantly in WT islets in response to the addition of glucose or KCl. This response was not attenuated in *Stard10KO* islets.

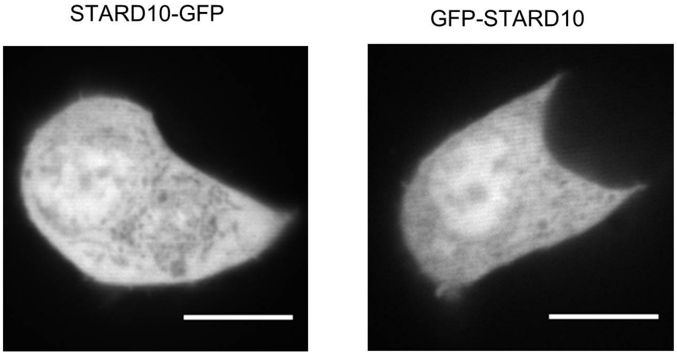


***Supplemental Figure 4. Subcellular localisation of STARD10-GFP (left panel) and GFP-STARD10 (right panel) in EndoC-βH1 cells***. Scale bar = 10µm.

***Supplemental Table 1. Characteristics of the human donors used for the electron microscopy analysis of insulin granules***

ND: normoglycemic donor; IGT: donor with impaired glucose tolerance; T2D: donor with type 2 diabetes; T3D: donor with type 3 diabetes

| **Gender** | **Diabetes status** | **STARD10 expression (Counts per million reads)** |
|---|---|---|
| F | T2D | 3.1216 |
| F | ND | 2.9139 |
| F | T2D | 2.8928 |
| F | T2D | 2.8561 |
| F | T2D | 2.6440 |
| M | T2D | 0.7689 |
| M | T3D | 0.1353 |
| F | IGT | 0.0206 |

***Supplemental Table 2. Enrichment of neuronal genes among the dowregulated genes in βStard10KO islets***

Genes identified by the GO consortium as components of "neuron projection" and "neuron part", found enriched among the downregulated genes identified in *βStard10*KO islets, listed as relative expression versus WT (log2 fold change).

| | symbol | log2FoldChange | pvalue | padj | entrezID |
|---|---|---|---|---|---|
| ENSMUSG00000029361 | Nos1 | -0.3533 | 4.34E-07 | 0.000468 | 18125 |
| ENSMUSG00000076441 | Ass1 | -0.3040 | 4.67E-05 | 0.014264 | 11898 |
| ENSMUSG00000030077 | Chl1 | -0.3032 | 1.91E-05 | 0.009003 | 12661 |
| ENSMUSG00000068748 | Ptprz1 | -0.2947 | 6.26E-06 | 0.00394 | 19283 |
| ENSMUSG00000029168 | Dpysl5 | -0.2805 | 0.00017 | 0.034308 | 65254 |
| ENSMUSG00000035864 | Syt1 | -0.2618 | 0.0002 | 0.038729 | 20979 |
| ENSMUSG00000029778 | Adcyap1r1 | -0.2588 | 8.67E-05 | 0.022575 | 11517 |
| ENSMUSG00000061576 | Dpp6 | -0.2531 | 2.89E-05 | 0.011209 | 13483 |
| ENSMUSG00000038077 | Kcna6 | -0.2514 | 0.000115 | 0.028134 | 16494 |
| ENSMUSG00000027500 | Stmn2 | -0.2474 | 3.69E-05 | 0.012594 | 20257 |
| ENSMUSG00000023945 | Slc5a7 | -0.2435 | 2.18E-05 | 0.009472 | 63993 |
| ENSMUSG00000033615 | Cplx1 | -0.2093 | 0.000111 | 0.027526 | 12889 |

***Supplemental Table 3. Gene ontology analysis of STARD10 binding partners in INS1 (832/13) cells: biological process***

Top 10 most significant gene ontology terms for biological processes

| GO biological process | Number of proteins | Fold Enrichment | P-value |
|---|---|---|---|
| RNA processing (GO:0006396) | 85 | 11.19 | 1.99E-62 |
| gene expression (GO:0010467) | 114 | 6.4 | 2.66E-62 |
| RNA splicing (GO:0008380) | 63 | 22.47 | 5.73E-62 |
| mRNA processing (GO:0006397) | 67 | 18.66 | 2.16E-61 |
| mRNA metabolic process (GO:0016071) | 71 | 14.76 | 6.70E-59 |
| RNA metabolic process (GO:0016070) | 93 | 7.16 | 5.13E-53 |
| cellular nitrogen compound metabolic process (GO:0034641) | 123 | 4 | 1.83E-46 |
| nucleic acid metabolic process (GO:0090304) | 96 | 5.17 | 3.55E-43 |
| mRNA splicing, via spliceosome (GO:0000398) | 43 | 21.42 | 3.54E-41 |
| RNA splicing, via transesterification reactions with bulged adenosine as nucleophile (GO:0000377) | 43 | 21.42 | 3.54E-41 |

***Supplemental Table 4. Gene ontology analysis of STARD10 binding partners in INS1 (832/13) cells: cellular components***

Top 10 most significant gene ontology terms for cellular components

| GO cellular component | Number of proteins | Fold Enrichment | P-value |
|---|---|---|---|
| ribonucleoprotein complex (GO:1990904) | 100 | 8.18 | 1.04E-62 |
| spliceosomal complex (GO:0005681) | 52 | 28.97 | 9.68E-56 |
| catalytic step 2 spliceosome (GO:0071013) | 40 | 41.38 | 7.06E-48 |
| protein-containing complex (GO:0032991) | 159 | 2.72 | 1.45E-43 |
| intracellular organelle part (GO:0044446) | 179 | 2.19 | 4.59E-39 |
| nuclear speck (GO:0016607) | 48 | 13.14 | 3.93E-37 |
| organelle part (GO:0044422) | 180 | 2.11 | 4.48E-37 |
| nuclear part (GO:0044428) | 128 | 3.04 | 2.57E-36 |
| U2-type spliceosomal complex (GO:0005684) | 31 | 36.48 | 8.32E-36 |
| nucleus (GO:0005634) | 159 | 2.36 | 1.90E-35 |

***Supplemental Table 5. Gene ontology analysis of STARD10 binding partners in INS1 (832/13) cells: molecular functions***

Top 10 most significant gene ontology terms for molecular functions

| GO molecular function | Number of proteins | Fold Enrichment | P-value |
|---|---|---|---|
| RNA binding (GO:0003723) | 93 | 7.99 | 6.96E-57 |
| nucleic acid binding (GO:0003676) | 112 | 3.33 | 6.80E-34 |
| mRNA binding (GO:0003729) | 39 | 13.91 | 5.38E-31 |
| heterocyclic compound binding (GO:1901363) | 132 | 2.45 | 5.21E-28 |
| organic cyclic compound binding (GO:0097159) | 132 | 2.4 | 3.06E-27 |
| binding (GO:0005488) | 184 | 1.43 | 1.51E-14 |
| mRNA 3'-UTR binding (GO:0003730) | 14 | 16.9 | 8.21E-13 |
| ATP-dependent RNA helicase activity (GO:0004004) | 11 | 28.77 | 1.70E-12 |
| RNA helicase activity (GO:0003724) | 11 | 27.99 | 2.18E-12 |
| RNA-dependent ATPase activity (GO:0008186) | 11 | 27.99 | 2.18E-12 |

*Supplemental Table 6. Statistics from STARD10 crystallographic analysis*

| **Complex** | **StarD10** |
|---|---|
| PDB ID | 6SER |
| **Data collection** | |
| Source | I03 |
| Wavelength (Å) | 0.9762 |
| Resolution (Å) | 60.06-2.30 (2.34-2.30) |
| Space group | I 2 2 2 |
| Cell dimensions: a, b, c | 68.3, 101.2, 125.8 |
| Observations | 256184 (11210) |
| Unique reflections | 19758 (925) |
| $R_{merge}$ (%) | 8.5 (44.0) |
| I/σI | 19.2 (5.6) |
| Completeness (%) | 99.9 (97.7) |
| Redundancy | 13.0 |
| **Refinement statistics** | |
| Resolution (Å) | 60.06-2.30 (2.34-2.30) |
| $R_{factor}$ (%)/$R_{free}$ (%) | 20.98/24.77 |
| rmsd bonds (Å)/angles (°) | 0.007/1.0 |
| Ramachandran plot: Favored (%) | 96.0 |

The values in parentheses refer to the last shell.
$R_{factor} = \Sigma||F(obs)- F(calc)||/\Sigma|F(obs)|$.
$R_{free}$ = R factor calculated using 5.0% of the reflection data randomly chosen and omitted from the start of refinement.
